# Luminal epithelial cells integrate variable responses to aging into stereotypical changes that underlie breast cancer susceptibility

**DOI:** 10.1101/2022.09.22.509091

**Authors:** Rosalyn W. Sayaman, Masaru Miyano, Parijat Senapati, Arrianna Zirbes, Sundus Shalabi, Michael E. Todhunter, Victoria Seewaldt, Susan L. Neuhausen, Martha R. Stampfer, Dustin E. Schones, Mark A. LaBarge

**Author notes:** Correspondence (MAL), (RWS). These authors contributed equally (Rosalyn W. Sayaman, Masaru Miyano).

## Abstract

Effects from aging in single cells are unpredictable, whereas aging phenotypes at the organ- and tissue-levels tend to appear as stereotypical changes. The mammary epithelium is a bilayer of two major phenotypically and functionally distinct cell lineages, the luminal epithelial and myoepithelial cells. Mammary epithelia exhibit substantial stereotypical changes with age that merits attention because they are putative breast cancer-cells-of-origin. We hypothesize that effects from aging that impinge upon maintenance of lineage fidelity increases susceptibility to cancer initiation. We identified two models of age-dependent changes in gene expression, directional changes and increased variance, which contributed to genome-wide loss of lineage fidelity. Age-dependent variant responses were common to both lineages, whereas directional changes were almost exclusively detected in luminal epithelia and implicated downregulation of chromatin and genome organizers such as *SATB1*. Epithelial expression of gap junction protein *GJB6* increased with age, and modulation of *GJB6* expression in heterochronous co-cultures revealed that it provided a communication conduit from myoepithelial cells that drove directional change in luminal cells. Age-dependent luminal transcriptomes comprised a prominent signal detectable in bulk tissue during aging and transition into cancers. A machine learning classifier based on luminal-specific aging distinguished normal from cancer tissue and was predictive of breast cancer subtype. We speculate that luminal epithelia are the ultimate site of integration of the variant responses to aging in their surrounding tissue and that their emergent aging phenotype both endows cells with the ability to become cancer-cells-of-origin and embodies a biosensor that presages cancer susceptibility.

## Introduction

The stereotyped aging phenotypes exhibited by organisms, organs, and tissues represent the integration of accumulated, stochastically incurred damages to individual cells that result in commonly understood hallmarks of aging (López-Otín et al., 2013; Todhunter et al., 2018). Age-associated directional changes in transcriptomes of whole tissues are well documented (de Magalhães et al., 2009; Glass et al., 2013; Peters et al., 2015; Volkova et al., 2005). These directional molecular changes explain, at least in part, the noticeable phenotypic changes that accompany aging. However, while increased susceptibility to a plethora of diseases, including cancers is a prominent consequence of aging, the manifestation and onset of diseases vary between same-aged individuals. Indeed, variance between individuals arises in the contexts of tumors, diet, and aging (Bashkeel et al., 2019; de Jong et al., 2019; Slieker et al., 2016; Xie et al., 2011). We propose that the variability across individuals may itself be an important molecular phenotype of aging, and individuals with outlier expression profiles provide an avenue for understanding biological processes that explain the differential cancer-susceptibility of aged individuals.

The breast is an excellent model system for examining aging at the molecular and cellular levels because normal tissue across the adult lifespan is available from common cosmetic and prophylactic surgeries. Cultured pre-stasis human mammary epithelial cells (HMEC) supports growth of all lineages from women across the lifespan (Garbe et al., 2009; Labarge et al., 2013) and enable detailed and reproducible molecular studies of cancer progression (Stampfer et al., 2013). Well-established lineage-specific markers and cell-sorting protocols facilitate experimentation at lineage-specific resolution. Furthermore, breast tissue provides an ideal model for studying aging-associated cancer susceptibility as 82% of new breast cancers are diagnosed in women ≥50y (DeSantis et al., 2019). Directional changes in gene expression with age were reported in whole breast tissue, including changes associated with breast cancer biological processes (Lee & Lee, 2017; Yau et al., 2007). However, aging is also associated with significant shifts in proportions of breast cell lineages, including epithelial and stromal populations (Benz, 2008; Garbe et al., 2012), so it is unclear how tissue-level molecular changes in normal aging reflect changes in cell-intrinsic and microenvironment states. Lineage-specific analyses are necessary to unravel such mechanisms.

The mammary epithelium, the origin of breast carcinomas, is a bilayer of two major phenotypically and functionally distinct cell lineages. Myoepithelial cells (MEPs) are basally located, contractile and have tumor suppressive properties (Pandey et al., 2010). Luminal epithelial cells (LEPs) are apically-located, secretory and include subpopulations of hormone receptor positive cells (Booth & Smith, 2006). We previously demonstrated loss of lineage fidelity as an aging phenomenon – where the faithfulness of expression of established lineage-specific markers diminishes with age without loss of the lineage-specificity of other canonical markers and the gross phenotypic and histological differences between LEPs and MEPs (Miyano et al., 2017). We hypothesize that the aging mechanisms that impinge upon the genome-wide maintenance of lineage fidelity are drivers of susceptibility to cancer initiation in breast tissue.

Here we demonstrate how age-dependent directional and variant transcriptional responses integrate in breast epithelia and explain how these changes could lead to increased susceptibility to cancer initiation. Through transcriptomic profiling of primary LEPs and MEPs we showed that loss of lineage fidelity in gene expression with age was a genome-wide phenomenon. We identified two models mediating loss of lineage fidelity in breast epithelia with age: (i) via directional changes identified through differential expression (DE); or (ii) via an increase in variances identified through differential variability (DV) analysis. Age-dependent DE explained part of the observed loss of lineage fidelity, while our model of the overall increase in variances with age also accounted for a comparable fraction of this loss. Directional changes in expression with age strikingly occurred almost exclusively in luminal cells, whereas changes in variance were found in both epithelial lineages. The genome-wide directional changes in LEPs involved dysregulation of chromatin and genome organizers such as *SATB1* with age, which we also detected in bulk tissue. Loss of lineage fidelity led to enrichment of genes and biological processes commonly dysregulated in cancers, and alteration of the LEP-MEP interactome that was significantly modulated by apical cell-cell junction proteins, such as *GJB6*. Modulation with shRNA *GJB6* in MEPs was sufficient to reduce the rate of molecular aging of adjacent LEPs as determined with a breast specific biological clock. Using machine learning, we showed that genes with age-dependent directional and variable changes in normal LEPs had predictive value in distinguishing normal breast tissue from breast cancers and classifying breast cancer PAM50 subtypes. Age-dependent changes in LEPs reflected dysregulation of biological processes that are convergent with breast cancer. The degree and variability of age-dependent changes across individuals may explain the differential susceptibility of specific individuals to breast cancer initiation, and to the development of specific breast cancer subtypes.

## Results

### Genome-wide loss of lineage-specific expression in breast epithelia with age

LEPs and MEPs were enriched from 4th passage finite-lifespan HMEC from reduction mammoplasties from two age cohorts: younger <30y women considered to be premenopausal (m_LEP_=16, m_MEP_=16, samples, n=11 subjects, age range 16-29y) and older >55y women considered to be postmenopausal (m_LEP_=11, m_MEP_=11, n=8, age range 56-72y) (**Figure 1—table supplement 1**). We analyzed the expression of 17,328 genes with comparable dynamic ranges and consistent lineage-specific expression between primary organoid and 4th passage LEPs and MEPs in both age cohorts (linear regression R^2^=0.88 to 0.91, *p*<0.0001) (**Figure 1—figure supplement 1A-1D**). To understand how lineage fidelity of the two epithelial cell types (**Figure 1A**) changes with age, we performed differential expression (DE) analysis comparing LEP and MEP expression in younger <30Y and older >55y women. DE genes between LEPs and MEPs decreased with age (adj. *p<*0.05, <0.01, <0.001) (**Figure 1—figure supplement 1E-1F**). Restricting analysis to genes with strong lineage-specific bias (DE adj*. p<*0.001, lfc≥1), we found 4,040 genes (23% of all genes analyzed) with highly significant lineage-specific DE in younger women– of which 59% were LEP-specific and 41% were MEP-specific. In contrast, 3,345 genes had highly lineage-specific DE in older women – of which 56% were LEP-specific and 44% were MEP-specific. Shifts in lineage-specific expression with age were illustrated in the strata-plot in (**Figure 1B**). Loss of lineage-specific expression with age occur genome-wide and was detected in 1,022 genes – a majority of which (65.5%) were LEP-specific.

**Figure 1.**
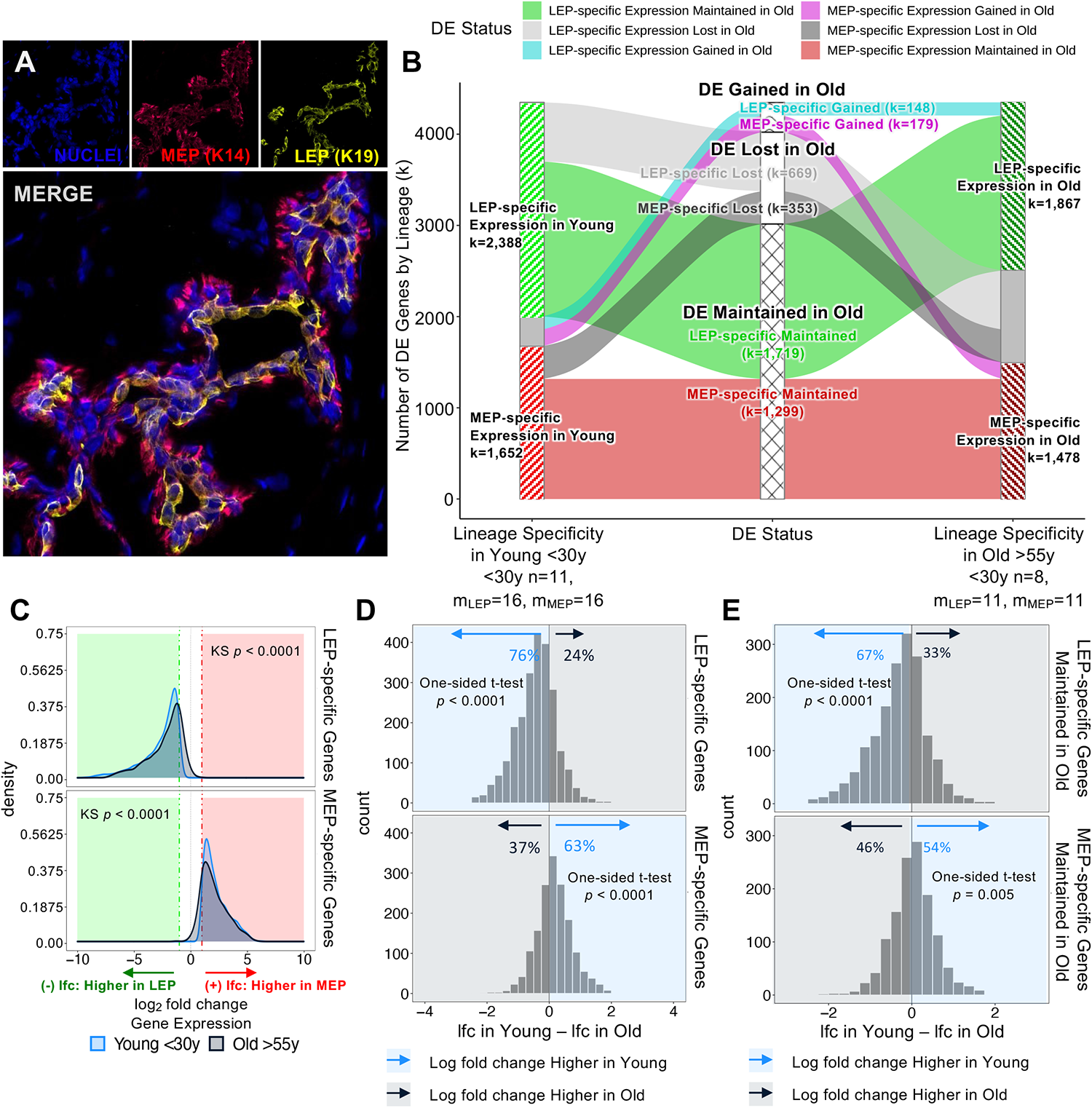
Genome-wide loss of lineage-specific expression with age. (**A**) Immunofluorescence staining of normal breast tissue showing the mammary epithelium with an apical LEPs (K19) surrounded by basal MEPs (K14). (**B**) DE LEP-specific and MEP-specific genes (adj*. p*<0.001, lfc≥1) in younger <30y (left) and older >55y (right) women. Strata plot shows changes in lineage-specific DE with age, showing the number of LEP- and MEP-specific genes gained (cyan and magenta), lost (light and dark gray), and maintained (green and red) in older women. Number of subjects (n) and sample replicates (m) in each DE analysis annotated; number of DE genes (k) in each age group and DE status indicated. (**C**) Distribution of lfc in expression between LEPs and MEPs in younger and older subjects for either DE LEP-specific (top panel) or MEP-specific (bottom panel) genes. KS *p*-values for equality of distributions of lfc between younger and older women annotated. (**D-E**) Histogram of pairwise differences in lfc in expression between LEPs and MEPs in younger vs. older women for (**D**) all genes with lineage-specific expression in younger women or (**E**) only genes that maintain lineage-specific expression in older women. Genes with LEP-specific and MEP-specific expression are shown in the top and bottom panels respectively. The percent of genes with higher lfc in younger women (light blue) or higher lfc in older women (blue gray) are indicated. One-sided t-test p-values annotated.

Lineage fidelity is the loss of the faithful expression of lineage-specific markers with age. Statistically, we described this loss as a phenomenon whereby the magnitude of gene expression differences that distinguish LEPs from MEPs decreased with age, which is seen as shifts in distributions of fold changes between lineages to smaller values in the older cohort (Kolmogorov-Smirnov two-sample test, KS *p<*0.0001) (**Figure 1C**). We found that 76% of LEP-specific genes and 63% of MEP-specific genes had higher fold changes between lineages in younger cells compared to older cells (**Figure 1D**). These percentages indicated loss of lineage fidelity was not restricted to genes that lost lineage-specific expression. Indeed, within the subset of genes where lineage-specific DE was maintained with age by significance threshold, the majority – 67% of LEP-specific genes and 54% of MEP-specific genes, still showed larger fold differences between LEPs and MEPs in younger women (**Figure 1E**). These data expand on our earlier findings that identified loss of lineage fidelity in a limited set of lineage-specific probes (Miyano et al., 2017), and underscore the genome-wide nature of this phenomenon whereby gene expression differences that distinguish the major epithelial lineages of the breast decrease with age.

### Loss of lineage fidelity with age leads to disrupted lineage-specific signaling

Because loss of lineage-specific expression could upset the relative balance of ligands and receptors in each lineage, we explored how loss of lineage fidelity could lead to disrupted or dysregulated cell-cell communication between neighboring cell types. We defined the breast interactome as a set of possible ligand-receptor interactions between cell populations based on the DE of cell-specific ligands and their cognate receptors in younger women. Using published ligand-receptor pairs (LRPs) (Ramilowski et al., 2015), we identified 224 candidate lineage-specific LRPs in younger LEPs and MEPs based on the DE of 62 LEP-specific and 66 MEP-specific ligands, and 45 LEP-specific and 47 MEP-specific cognate receptors (**Figure 2—figure supplement 1A**). Protein-protein interaction (PPI) functional enrichment of lineage-specific LRPs identified top KEGG canonical biological pathways (FDR *p<*0.001) (**Figure 2—figure supplement 1B-1C**), with ligands and receptors related to cytokine-cytokine receptor interaction, PI3K-Akt, MAPK and Rap1 signaling commonly enriched in LEPs and MEPs. Enrichment of cytokine, immune and infection-related pathways further suggested lineage-specific interactions between epithelial and immune cells. LEP-specific LRPs were enriched for cell adhesion molecules (CAMs) involved in cell-cell and cell-ECM interactions, and axon guidance molecules (AGMs), while MEP-specific LRPs were enriched for ECM-receptor interaction and focal adhesion LRPs.

Loss of lineage fidelity with age led to disruption of 74 LRPs based on the loss of lineage-specific expression of ligands and/or their cognate receptors (**Figure 2A**). For each lineage, we considered KEGG canonical biological pathways (FDR *p<*0.01) that were likely to exhibit dysregulated signaling either through direct disruption of the LRPs due to loss of lineage-specific signaling of the ligand or receptor, or indirect loss of signaling homeostasis via dysregulation of its cognate pair (**Figure 2—figure supplement 1B-1C**). Loss of lineage-specific expression of LEP LRPs with age was enriched for canonical pathways involved in (1) cell-cell and cell-ECM interactions including CAMs, AGMs, and adherens junctions, and (2) cytokine, immune and infection-related pathways. Loss of lineage-specific expression of MEP LRPs with age were associated with (1) pathways in cancer; (2) pathways involved with MAPK, EGFR, NOTCH and PI3K-AKT signaling; and (3) MEP contractility. These findings suggest that loss of lineage fidelity with age has the potential to affect a wide range of biological processes regulating lineage-specific function and signaling, including potential dysregulation of cancer-related processes and immune-specific signaling by the epithelia.

**Figure 2.**
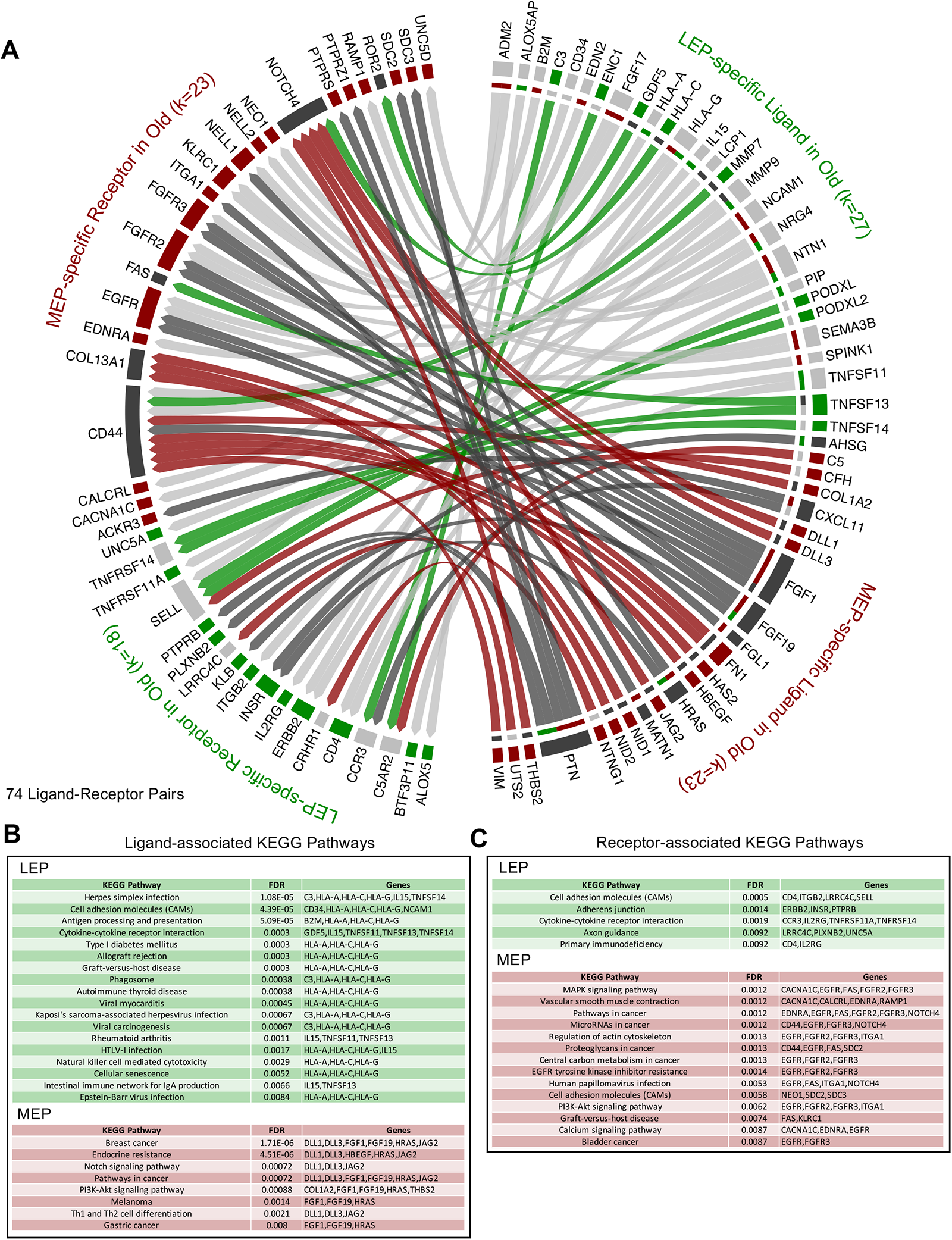
Loss of lineage fidelity with age leads to disrupted lineage-specific signaling. (**A**) Interactome map of DE lineage-specific ligand-receptor pairs (LRPs) (adj*. p*<0.001, lfc≥1) that show loss of lineage-specific expression of either ligands and/or their cognate receptors in older LEPs (light gray) or MEPs (dark gray). LRPs are connected by chord diagrams from the cell type expressing the ligand (L) to the cell type expressing the cognate receptor (R). Number of LRPs, and genes (k) in each category annotated. (**B-C**) Network functional enrichment of top KEGG pathways (FDR *p*<0.001) associated with loss of lineage-specific DE of (**b**) ligands and/or (**c**) cognate receptors in LEPs and MEPs in older women.

### Models of loss of lineage fidelity in breast epithelia

To understand the changes within each cell population that contribute to the observed aging-associated loss of lineage fidelity, we explored two models that could explain the decrease in DE between LEPs and MEPs with age. The first model took into account age-dependent directional changes either through stereotypic, up- or down-regulation, leading to loss of lineage-specific expression – e.g., LEPs acquire MEP-like expression patterns and/or MEPs acquire LEP-like expression patterns in the older cohort (**Figure 3Ai**). The second model considered aging-associated increase in variances in the expression of lineage-specific genes in LEPs and/or MEPs from older women, leading to a loss of detection of DE between lineages (**Figure 3Aii**). We describe the contributions of each in the following sections.

### The luminal lineage is a hotspot for age-dependent directional changes

There was an extreme lineage bias in the numbers of DE genes between younger and older cells, with the majority of age-dependent changes occurring in LEPs. In LEPs, 471 genes were DE as a function of age; in contrast, in MEPs only 29 genes were DE with age (adj. *p<*0.05) (**Figure 3B**). Moreover, we identified age-dependent changes that showed lineage independence with five genes: *LRRC4, PSORS1C1, SCNN1B, ZNF518B,* and *ZNF521* commonly DE across cell types, leaving only 24 genes changed with age exclusively in MEPs. That stereotypic directional changes associated with aging were almost exclusively found in LEPs suggests that this lineage could serve as a primary indicator of aging – a proverbial canary in the coalmine.

**Figure 3.**
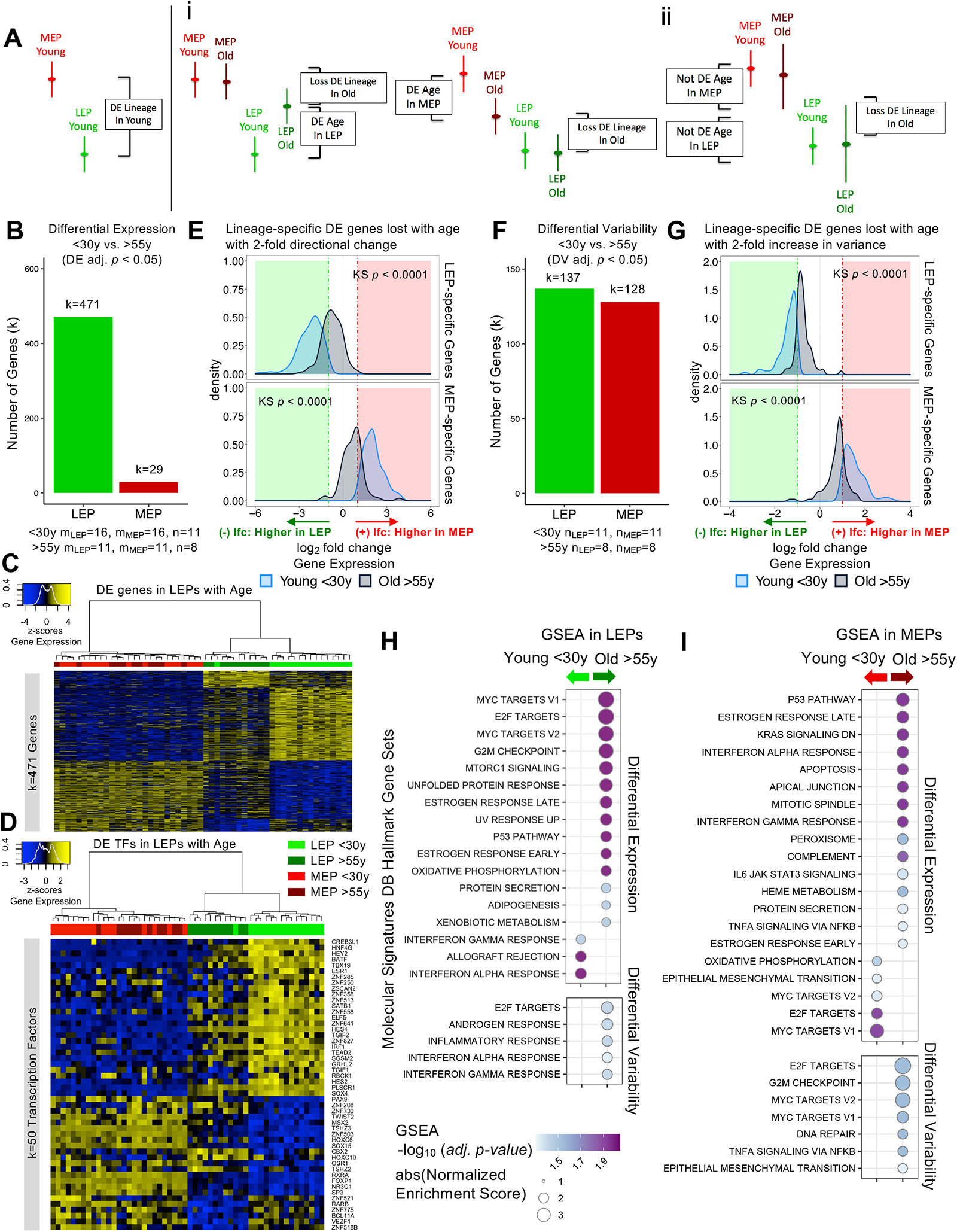
The luminal lineage is a hotspot for age-dependent directional changes. (**A**) Models of loss of lineage fidelity illustrate hypothesized mechanisms leading to loss of lineage fidelity: (i) Age-dependent DE shifts in gene expression in LEPs and/or MEPs of older relative to younger cells; or (ii) An increase in gene expression variance in older LEPs and/or MEPs that lead to loss of detection of lineage-specific DE between LEPs and MEPs with age. (**B**) Number of DE genes (adj*. p<*0.05) between younger and older LEPs or MEPs. Number of subjects (n) and sample replicates (m) in DE analysis annotated. (**C-D**) Hierarchical clustering of all LEP and MEP samples based on sample-level expression of age-dependent (**C**) DE genes in LEPs (adj. *p* < 0.05) and (**D**) DE transcription factors in LEPs (adj. *p* < 0.05). Number of DE genes (k) indicated. Gene expression scaled regularized log (rlog) values are represented in the heatmap; clustering performed using Euclidean distances and Ward agglomerative method. (**E**) Distribution of lfc in expression between LEPs and MEPs in younger and older women for LEP-specific (top panel) or MEP-specific (bottom panel) genes that are lost with age (DE adj*. p*<0.001, lfc ≥1) and that have at least a 2-fold age-dependent directional change in the older cohort. KS *p*-values annotated. (**F**) Number of DV genes (adj*. p<*0.05) between younger and older LEPs or MEPs. Number of subjects (n) in DV analysis annotated. (**G**) Distribution of lfc in expression between LEPs and MEPs in younger and older women for LEP-specific (top panel) or MEP-specific (bottom panel) genes that are lost with age (DE adj*. p* <0.001, lfc≥1) and that have at least a 2-fold age-dependent increase in variance in the older cohort. KS *p*-values annotated. (**H-I**) MSigDB Hallmark gene sets identified by GSEA to be enriched (adj*. p<0.05*) in younger and older (**H**) LEPs and (**I**) MEPs based on age-dependent DE (top) or DV (bottom).

Age-dependent differential upregulation (251 genes) and downregulation (220 genes) of LEP gene expression (adj. *p<*0.05) occurred at comparable frequencies (**Figure 3C**). In LEPs, 82% of the genes that were DE changed in a direction towards acquiring MEP-like patterns with age (**Figure 3C**). Although changes in MEPs were far fewer, we note that shifts in expression and methylation in older MEPs led to LEP-like patterns (**Figure 3—figure supplement 1A**).

Because dysregulation of regulatory factors like transcription factors (TFs) could lead to further dysregulation of downstream targets, we compared TF expression between younger and older cells in each lineage. Expression of key TFs (Lambert et al., 2018) were significantly altered in older cells, with 50 TFs showing age-dependent DE in LEPs, and 4 TFs DE in MEPs (adj*. p<*0.05), the majority of which have known roles in breast cancer progression. Of these DE TFs in LEPs, 88% changed expression in older LEPs towards the direction of MEP-like expression (**Figure 3D**). These included highly expressed TFs in younger LEPs that were down-regulated with age, such as: LEP-specific TF *ELF5* (Miyano et al., 2021), *GRHL2*, *SGSM2*, *HES4*, *ZNF827*, and genome organizer *SATB1* (**Figure 3—figure supplement 1Bi-v**). Loss of *GRHL2* and *SGSM2* are associated with down-regulation of E-cadherin and epithelial-to-mesenchymal transition (EMT) in mammary epithelial cells (Lin et al., 2019; Xiang et al., 2012). *HES4* is a canonical target gene of Notch1 which plays an important role in normal breast epithelial differentiation and cancer development (Kontomanolis et al., 2018). *ZNF827* mediates telomere homeostasis through recruitment of DNA repair proteins (Vilas et al., 2018). And *SATB1* has genome organizing functions in stem cells and tumor progression (Kohwi-Shigematsu et al., 2013). Several TFs also gained expression in older LEPs, including: *SP3* and *ZNF503* (**Figure 3—figure supplement 1Bvi-vii**). *SP3* silencing inhibits Akt signaling and breast cancer cell migration and invasion (Mansour, 2020). *ZNF503* inhibits *GATA3* expression, a key regulator of mammary LEP differentiation, and down regulation is associated with aggressive breast cancers (Kouros-Mehr et al., 2006; Shahi et al., 2017). Age-dependent dysregulation of TFs in LEPs may presage larger-scale changes through TF binding of gene regulatory regions of downstream targets in older LEPs.

To understand the contribution of age-dependent directional changes in driving loss of lineage fidelity (**Figure 3Ai**), we examined the overlap of age-dependent DE genes with genes that lost lineage-specific DE. Only 9% of the loss in lineage-specific DE was explained by age-dependent DE in LEPs or MEPs at adj. *p<*0.05. If we considered all genes with at least 2-fold change DE with age, these age-dependent directional changes accounted for only 21% of loss of lineage-specific expression events, leading to a significant decrease in the magnitude of expression fold changes between LEP- and MEP-specific expression in the older cells (**Figure 3E**). These findings suggest that other mechanisms play a substantial role in regulating lineage fidelity, and that molecular changes associated with aging are not limited to stereotyped directional changes.

### Aging-associated increase in variance contributes to loss of lineage fidelity

Next, we explored the alternate model that incorporated measures of variance as an explanation for the loss of lineage-specific expression in older epithelia (**Figure 3Aii**). Gene expression means and variances (**Figure 3—figure supplement 1C-1F**) of LEPs and MEPs from younger cells were categorized into quantiles and corresponding categories in older cells were then assessed. Gene expression means shifted minimally between younger and older cells (**Figure 3—figure supplement 1C-1D**), whereas shifts in variances occurred at a much higher frequency (**Figure 3—figure supplement 1E-1F**). Though the dynamic ranges of gene expression in LEPs and MEPs changed as a function of age, these changes were not stereotyped across individuals – i.e., different aged individuals had different sets of genes that deviated from the range of expression seen in younger samples.

Differential variability (DV) analysis identified 137 genes in LEPs and 128 genes in MEPs with significant age-dependent DV (adj. *p<*0.05) (**Figure 3F**). Twelve regulatory TFs in either LEPs or MEPs that had tuned windows of expression in younger cells were dysregulated in older cells through a significant increase in variance (adj. *p<*0.05) (**Figure 3—figure supplement 1G-1H**) and included *HES4, GLI1* and *KDM2B* in LEPs, and *HES6* in MEPs. *HES4*, which was also DE with age, is a known Notch target. *GLI1* activates the hedgehog pathway in mammary stem cells (Bhateja et al., 2019). Estrogen-regulated *HES6* is known to enhance breast cancer cell proliferation (Hartman et al., 2009). And lastly, *KDM2B (FBXL10)*, which was down-regulated in a subset of older women (**Figure 3—figure supplement 1Bviii**), is a histone demethylase ZF-CxxC protein that binds unmethylated DNA. These analyses suggested that age-dependent variability in expression across individuals can lead to differential outcomes as different downstream targets could be modulated in different individuals.

To understand how our model of age-dependent variability affected lineage-specific expression, we focused on genes that lost lineage-specific expression with age and that showed at least a 2-fold increase in variance in the older cohort (**Figure 3G**). Genes with 2-fold increases in variances with age explained 27% of the observed loss of lineage-specific expression events, on a par with the proportion (21%) explained by genes that had 2-fold changes in DE. Both of our models of directional and variant changes with age led to a significant decrease in the differential magnitude of LEP- and MEP-specific expression in the older cells (**Figure 3E, 3G**).

These data suggest that increased variances in transcription are considerable drivers of the loss of lineage fidelity in breast epithelia. The observed variances across the older cohort may underlie the age-dependent dysregulation of susceptibility-associated biological processes in specific individuals.

### Hallmark pathways associated with cancer are dysregulated with age in luminal and myoepithelial lineages

Gene set enrichment analysis (GSEA) identified Molecular Signatures Database (MSigDB) hallmark gene sets that were dysregulated with age, including gene sets known to be dysregulated in breast cancers that were enriched in older LEPs and MEPs (**Figure 3H-3I**).

Seventeen hallmark gene sets were significantly modulated in LEPs (adj*. p<*0.05) based on DE (**Figure 3H** top). Three immune-related gene sets were enriched in younger LEPs and included genes up-regulated in response to interferon IFN-alpha and -gamma, and during allograft rejection. In contrast, 14 gene sets were enriched in older LEPs, which included: genes regulated by MYC; genes encoding cell-cycle related targets of E2F TFs and involved in the G2/M checkpoint; genes upregulated by mTORC1 complex activation and during unfolded protein response; and genes involved in the p53 and protein secretion pathways.

Twenty hallmark gene sets were significantly modulated in MEPs (adj*. p<*0.05) based on DE (**Figure 3I** top). Five gene sets were enriched in younger MEPs, including: MYC and E2F targets; and genes defining EMT. In contrast, 15 gene sets were enriched in older MEPs, including: genes involved in p53 pathways; genes down-regulated by KRAS activation; genes mediating programmed cell death by caspase activation (apoptosis); immune-related gene sets upregulated in response to IFN-alpha, IFN-gamma and by IL-6 via STAT3, genes regulated by NF-kB in response to TNF and genes encoding components of the innate complement system; and genes encoding components of apical junction complex.

Five gene sets were significantly modulated (adj*. p<*0.05) based on DV and were enriched in older LEPs (**Figure 3H** bottom). These included: E2F targets that were similarly enriched via DE; genes defining responses to inflammation; and genes upregulated in response to IFN-alpha and IFN-gamma – gene sets that in contrast were enriched via DE in LEPs of younger women. Seven gene sets were significantly modulated (adj*. p<*0.05) based on DV and enriched in older MEPs (**Figure 3I** bottom). These included: genes involved in DNA repair and G2/M checkpoint; genes regulated by NF-kB in response to TNF – a gene set similarly enriched via DE; as well as MYC targets, E2F targets, and genes defining EMT – gene sets that in contrast were enriched via DE in MEPs from younger women.

Several enriched gene sets were involved in processes that were disrupted with age either via DE or DV in LEPs and MEPs, and such overlaps likely suggest integration of directional and variant responses and reflect their convergent impact in common biological processes. Furthermore, the divergence in the age-dependent DE and DV enrichment of cellular processes, such as MYC gene targets and genes involved in immunomodulatory signaling, suggests the genes that become variable with age are associated with pathways that are otherwise important in maintaining lineage-specificity and -function in younger cells.

### Age-dependent directional changes in the luminal lineage are indicators of aging breast tissue

Because LEPs dominated the age-specific signal amongst epithelia, we examined if the age-dependent DE contribution of the luminal lineage was detectable in bulk normal primary breast tissue (GSE102088, n=114) (Song et al., 2017). Genome-wide analysis identified 97 genes to be DE between younger <30Y and older >55y tissues (adj*. p<*0.05), the relatively smaller number of genes compared to age-dependent DE observed in LEPs likely due the cellular heterogeneity found in bulk tissue. To characterize the contribution of the LEP lineage to aging biology of the breast, we next performed gene set enrichment analysis (GSEA) to assess enrichment of LEP-specific age-dependent DE genes at tissue level. We found significant enrichment of differentially upregulated genes identified in young <30y LEPs in tissue from younger women (adj. *p*=0.012) (**Figure 4A**) and differentially upregulated genes identified in old >55y LEPs in tissue from older women (adj. *p*=0.006) (**Figure 4B**). These GSEA results indicate that while age-dependent changes in other cell populations may confound detection of the LEP-specific signal, age-dependent changes in LEPs were still reflected in bulk tissue.

**Figure 4.**
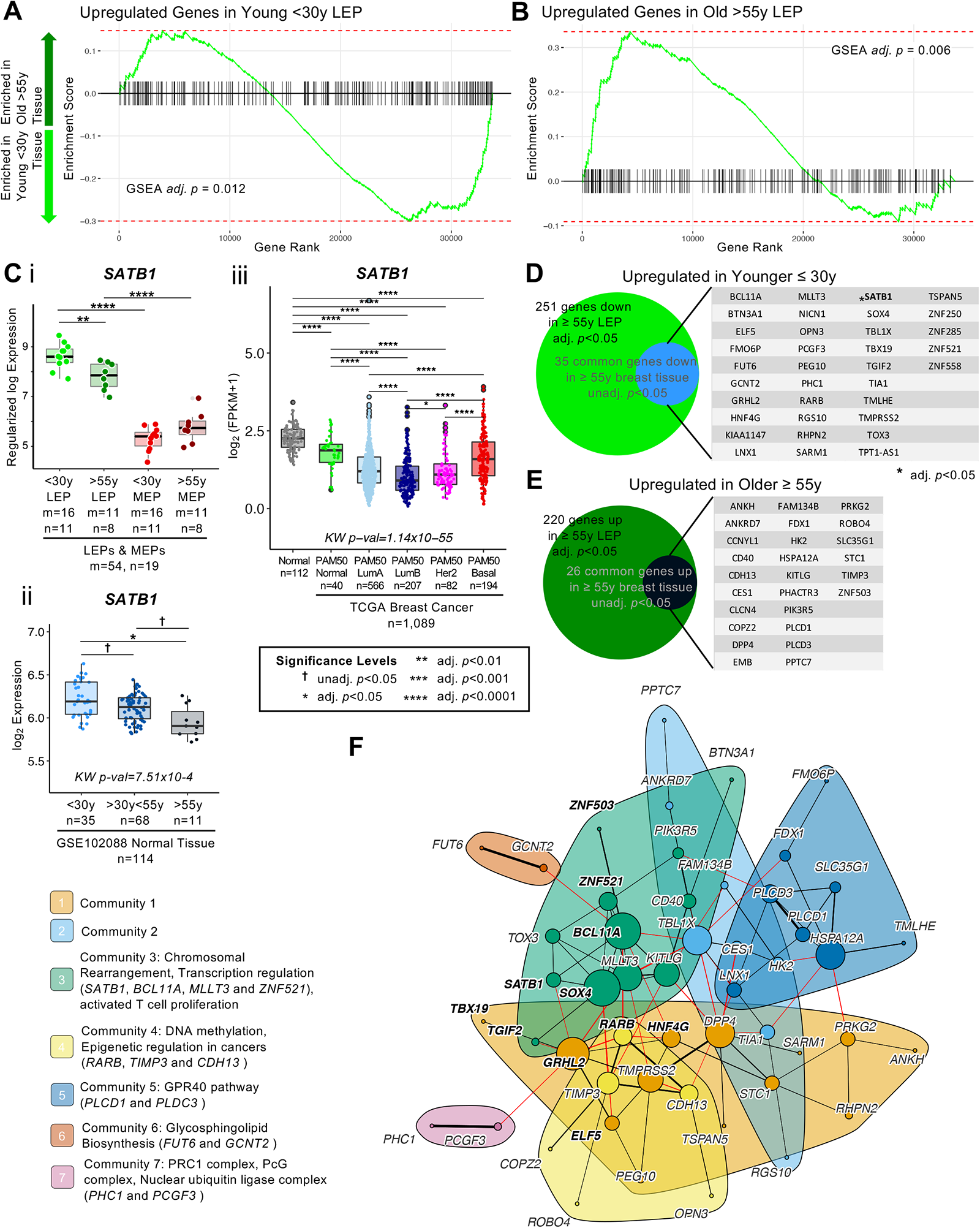
Age-dependent directional changes in the luminal lineage are indicators of aging breast tissue. (**A-B**) GSEA enrichments plots from age-dependent DE analysis of bulk tissue showing gene ranks based on DE test statistics and gene set enrichment scores. Age-dependent enrichment of two gene sets composed of (**A**) differentially upregulated genes in younger <30y LEPs and (**B**) differentially upregulated genes in older >55y LEPs in bulk tissue are shown. Negative enrichment scores indicate upregulation of specified gene set in tissue from younger <30y women, while positive enrichment scores indicated upregulation in tissue from older >55y women. GSEA enrichment BH adj. *p*-values are annotated. (**C**) Boxplots of *SATB1* gene expression: (i) subject-level rlog values in LEPs and MEPs of younger and older women; (ii) log_2_ values in normal breast tissue (GSE102088); and (iii) log_2_ FPKM values in the TCGA breast cancer cohort by PAM50 subtype in cancers and in matched normal tissue. Age-dependent DE adj*. p*-values in normal breast tissue and LEPs and lineage-specific DE adj*. p*-values in LEPs and MEPs are indicated (i-ii). Pair-wise Wilcoxon *p*-values between groups, and KW *p*-value across matched normal and breast cancer subtypes in TCGA annotated (iii). Number of subjects (n) and sample replicates (m) in each analysis annotated. (**D-E**) Venn diagram of genes with age-dependent DE in LEPs (adj. *p*<0.05) and at least nominal DE (unadj. *p*<0.05) in normal primary breast tissue. Genes commonly (**D**) upregulated and (**E**) downregulated in LEPs and bulk tissue with age are listed. (**F**) PPI network of common age-dependent DE genes in LEPs (adj*. p<*0.05) and bulk tissue (unadj. *p*<0.05) with TFs annotated in bold. Seven gene communities identified; corresponding network functional enrichment (FDR *p*<0.05) of selected processes annotated.

We then explored the GSEA leading edge genes – genes with the largest contribution to the significant enrichment of the LEP-specific age-dependent genes in bulk tissue. Of the leading edge genes, we found genome organizer *SATB1*, which showed significant LEP-specific expression compared to MEPs (adj*. p<*0.001, lfc≥1) (**Figure 4Ci**), to have the strongest signal in bulk tissue (**Figure 4Cii**). *SATB1* was significantly downregulated in both LEPs and breast tissue of older relative to younger women (adj*. p<*0.05) (**Figure 4Ci-ii**). This decrease with age in *SATB1* was also detected in normal breast tissue of women with cancer in the TCGA cohort (n=111, Wilcoxon adj*. p<*0.001) (**Figure 4—figure supplement 1A**). In TCGA breast cancers (n=1,089), PAM50 Luminal A (LumA), Luminal B (LumB) and Her2-enriched (Her2) breast cancer subtypes had the lowest expression of *SATB1* relative to PAM50 Basal-like (Basal) and Normal-like (Normal) intrinsic subtypes (KW adj*. p<*0.0001) (**Figure 4Ciii**). Moreover, in primary tumors with matched normal tissue (n=114 tumor and n=109 normal), we found *SATB1* to be significantly downregulated in PAM50 LumA, LumB and Her2 breast cancers relative to their matched normal tissue (**Figure 4—figure supplement 1B**). Together, these results suggest that *SATB1*-mediated genome organization may play a regulatory role in the maintenance of the luminal lineage and the observed genome-wide dysregulation with age and breast cancer.

Since we expected the signal in bulk tissue to be muted due to cellular heterogeneity, we also explored leading edge genes with nominally significant DE between younger and older tissue (unadj. *p*<0.05). Of the 251 genes upregulated in younger LEPs, 35 genes (14%) show nominally significant differential upregulation in young tissue (**Figure 4D**), including EMT-associated *GRHL2* (**Figure 3—figure supplement 1Ai**) and LEP-specific TF *ELF5* which we had previously shown to be predictive of accelerated aging in genetically high risk LEPs (Miyano, Sayaman et al., 2021). Of the 220 genes upregulated in older LEPs, 26 genes (12%) show nominally significant differential upregulation in old tissue (**Figure 4E**), including GATA3 inhibitor *ZNF503* (**Figure 3—figure supplement 1Avii**). Of the 61 genes we identified to be commonly dysregulated between younger and older LEPs and breast tissue, 17 were LEP-specific and 14 were MEP-specific in our lineage-specific DE analysis.

Common age-dependent DE genes between LEPs and bulk tissue showed significant PPI network enrichment (PPI enrichment *p*=0.014), including a 51-gene network that involved 11 DE TFs (**Figure 4F**) and 10 genes with high connectivity in the network (degree >10) that are potential nodes of integration. These include genes downregulated in the older group: TF *BCL11A –* a subunit of the BAF (SWI/SNF) chromatin remodeling complex (Kadoch et al., 2013), TF *SOX4 –* involved in determination of cell fate, TF *GRHL2*, *MLLT3 –* a chromatin reader component of the super elongation complex (SEC) (Moustakim et al., 2018); and genes upregulated in the older group: *DPP4 (CD26) – a* cell surface receptor involved in the costimulatory signal essential T-cell activation (Ikushima et al., 2000), *HSPA12A –* heat shock protein associated with cellular senescence, and *KITLG* – a ligand for luminal progenitor marker c-KIT in breast (Kim & Villadsen, 2018). Community detection algorithm identified 7 communities (**Figure 4F**). Functional network enrichment (FDR<0.05) showed that Community 3 anchored by TFs *BCL11A* and *SOX4* was enriched for genes associated with transcriptional regulation. *SATB1*, *BCL11A*, *MLLT3* and *ZNF521* were linked to chromosomal rearrangement and were downregulated in LEPs and breast tissues of older women. These genes showed breast cancer subtype-specific expression, and *BCL11A* and *ZNF521* in particular were downregulated in PAM50 LumA, LumB and Her2 tumors relative to matched normal tissue (**Figure 4—figure supplement 2Ai-iii, 2Bi-iii**). Community 4 members *RARB*, *TIMP3* and *CDH13* have been implicated as tumor suppressor gene targets of DNA methylation and epigenetic regulation in cancers. In addition, community 7 members *PHC1* and *PCGF3* are components of the Polycomb group (PcG) multiprotein polycomb repressor complex (PRC)-PRC1-like complex which is required for maintenance of the transcriptionally repressive state of many genes throughout development. *PHC1* and *PCGF3* were downregulated in LEPs and breast tissues of older women and showed breast cancer subtype-specific expression, with *PCGF3* downregulated in PAM50 LumB and Her2 tumors relative to matched normal tissue (**Figure 4—figure supplement 2Aiv-v, 2Biv-v**).

Taken together, genes commonly DE in younger and older LEPs and breast tissue either reflect stereotypic aging-associated molecular changes across different breast cell populations or are driven by LEP-specific changes, suggesting that age-dependent molecular changes in LEPs contribute to essential processes involved in the aging biology of the entire breast, and that are dysregulated in cancers.

### Genes encoding for cell-cell junctional proteins are dysregulated in aging epithelia

We showed previously that MEPs can impose aging phenotypes on LEPs – with LEPs from younger women acquiring expression patterns of older LEPs when co-cultured on apical surfaces of MEPs from older women (Miyano et al., 2017). This non-cell autonomous mechanism of aging requires direct cell-cell contact between LEPs and MEPs, suggesting that cell-cell junctional proteins play a role in age-dependent dysregulation in LEP-MEP signaling. Indeed, we identified apical junction-associated genes to be significantly enriched with age in MEPs (**Figure 3I** top).

We explored age-dependent dysregulation of a curated set of genes encoding for membrane components of adherens junctions, tight junctions, gap junctions, desmosomes, and cell adhesion molecules (CAMs) in LEPs and MEPs to identify candidate genes that may regulate communication between the lineages. Because age-dependent changes involve both DE and DV, we performed the non-parametric Lepage test to jointly monitor the central tendency and variability of gene expression of 198 genes encoding for cell-surface junction proteins between the younger and older cohorts. We found 42 genes were modulated in LEPs and/or MEPs with age (Lepage test *p<*0.05) (**Figure 5—figure supplement 1A**). These include genes that were modulated via a significant directional change with age such as the desmosomal cadherins genes, *DSG3* (desmoglein) and *DSC3* (desmocollin), which have been previously shown to be expressed in both LEPs and MEPs (Garrod & Chidgey, 2008) (**Figure 5—figure supplement 1Bi-ii**); and the genes encoding for essential tight junction components, *CLDN10* and *CLDN11* (**Figure 5—figure supplement 1Biii-iv**).

Gap junction *GJB6* (Connexin-30), which is expressed in both LEPs and MEPs in the normal mammary gland and forms homo-(LEP-LEP) and hetero-cellular (LEP-MEP) channels (Teleki et al., 2014) (**Figure 5A**), is of specific interest as it showed modulation via an increase in variance in older MEPs (*p=*0.02) and nominal increase in variance in older LEPs (*p=*0.06) (**Figure 5B**). As such, modulation of *GJB6* provided an avenue for exploring changes that could occur in both lineages and in only a subset of older women that may thus lead to differential susceptibility across aged individuals. To understand the transcriptional regulation of the GJB6 junctional protein, we explored the ChIP-seq (Cistromics) mammary gland data from The Signaling Pathways Project (SPP) Ominer database (**Figure 5C**). Nine TFs had binding signals within +/-10 of the TSS of *GJB6*, including progesterone receptor PGR, MYC and the LEP-specific TF *ELF5* – which we have previously shown to be regulated via direct LEP-MEP interactions in co-culture studies (Miyano et al., 2017).

**Figure 5.**
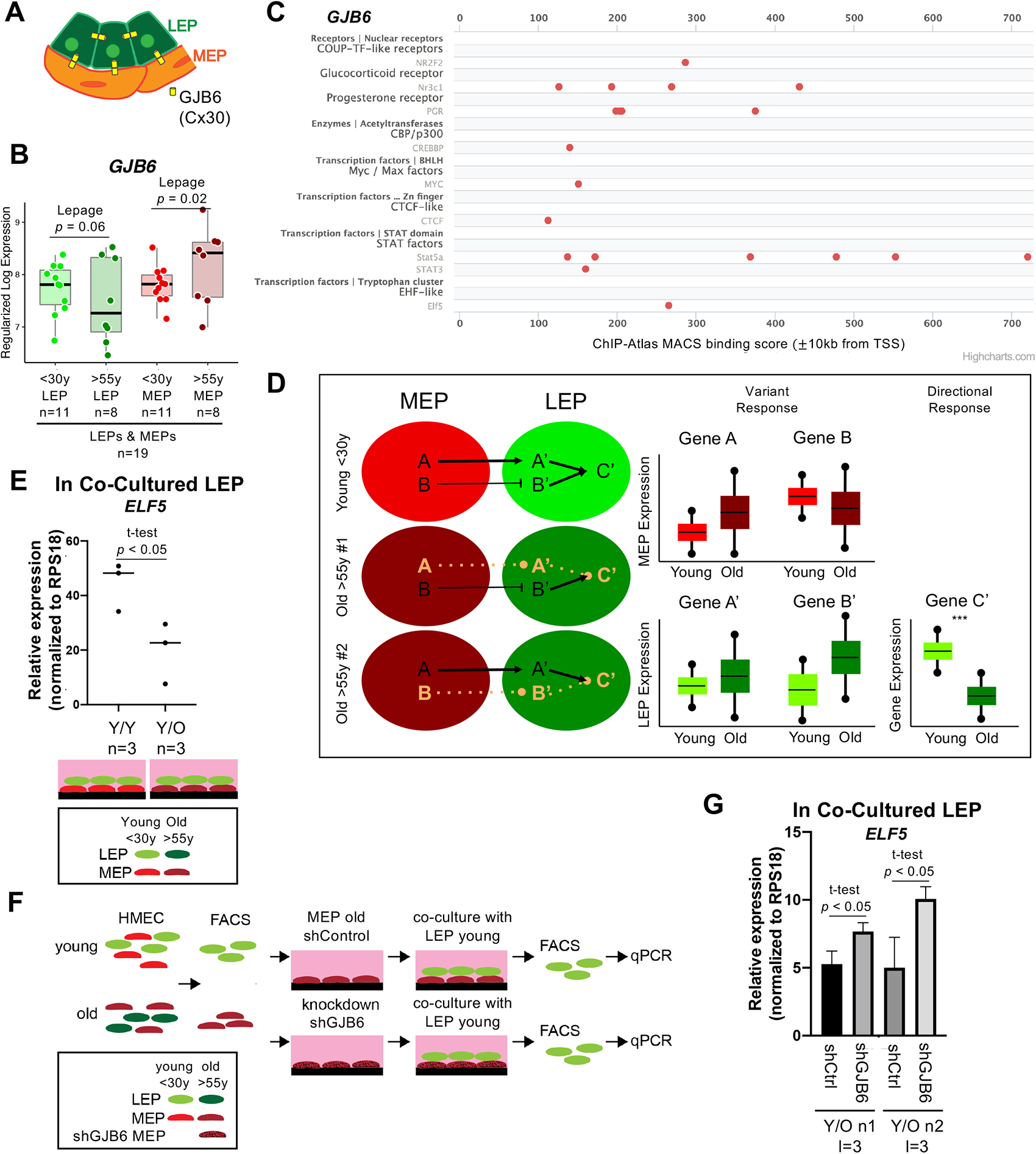
*GJB6* is a mediator of the non-cell autonomous mechanism of aging in breast. (**A**) Schematic of gap junction protein, Connexin-30 (*GJB6*) homotypic channel formation between LEPs and between LEP and MEP. (**B**) Boxplot of *GJB6* subject-level rlog expression values in LEPs and MEPs in younger and older women. Lepage test *p*-values are indicated. Number of subjects (n) in analysis annotated. (**C**) *GJB6* ChIP-seq (Cistromics) binding signal within +/-10 from the TSS in mammary gland (SPP Ominer database). (**D**) Schematic illustrating integration of directional and variant responses in older epithelial cells. Different genes are dysregulated in LEPs and MEPs of older individuals leading to an increase in variance in expression across aged cells. Through cell-cell signaling, variant responses in MEPs (gene A or gene B) lead to variant responses in LEPs (gene A’ or gene B’). Where these variant responses integrate and affect common downstream genes in LEPs (gene C’) lead to detectable age-dependent directional changes (***) that are seen as stereotyped responses in the lineage. (**E**) Relative expression of *ELF5* in younger LEPs co-cultured with either younger (Y/Y) or older (Y/O) MEPs. Two-tailed t-test *p*-value indicated. (**F**) Schematic of co-culture methodology with HMEC cells from younger and older women enriched by FACS for LEPs and MEPs; *GJB6* knock-down in older MEP feeder layer by shRNA; younger LEPs are co-cultured on top of the older MEP feeder layer for 10 days; LEPs separated from MEPs by FACS; and LEP expression levels measured by qPCR. (**G**) Relative expression of *ELF5* in younger LEPs co-cultured with either shControl or shGJB6 older MEPs. Two-tailed t-test *p*-value indicated. Number of subjects (n) and technical replicates (l) annotated.

### Gap Junction protein *GJB6* is a mediator of the non-cell autonomous mechanism of aging in breast

Because changes in MEPs are predominantly associated with DV rather than DE, we hypothesized that LEPs serve as integration nodes for dysregulation in MEPs where variant changes converge via common pathways leading to directional changes in genes downstream of these pathways (**Figure 5D**). We identified TF *ELF5* to be one such target; indeed, expression of the highly LEP-specific TF was dynamic and responsive to microenvironment changes (Miyano et al., 2017), and serves as an independent biological clock in breast (Miyano et al., 2021). *ELF5* was downregulated in younger LEPs when co-cultured on apical surfaces of older MEPs for 10 days (**Figure 5E**), concordant with the observed phenomenon of *ELF5* downregulation in LEPs with age.

We asked whether knockdown or inhibition of *GJB6* expression in the subset of older MEPs with higher expression relative to younger MEPs could restore proper signaling between LEPs and MEPs. To test this, we used our established heterochronous co-culture system and used recovery of LEP expression of *ELF5* as a readout (**Figure 5F**). If bringing variant *GJB6* under tighter control prevents chronologically older MEPs from imposing older biological ages in younger LEPs, then *ELF5* levels should not decrease in co-culture. FACs-enriched LEPs from younger women were co-cultured for 10 days on older MEPs treated with either shGJBG or scramble shRNA (shCtrl) (**Figure 5F, Figure 5—figure supplement 1C**). When co-cultured on top of older MEP-shGJB6 relative to MEP-shCtrl, LEP-expression of *ELF5* was maintained at higher levels (**Figure 5H**), consistent with higher expression levels in younger women. LEP-expression of *ELF5* likewise showed a stepwise (though non-significant) increase when older MEP feeder layers were pre-treated with increasing concentrations of a non-specific gap junction inhibitor 18 alpha-glycyrrhetinic acid (18αGA) (**Figure 5—figure supplement 1D**). Thus, reducing the level or variance of *GJB6* prevented older MEPs from imposing an older biological age in younger LEPs as determined by *ELF5* expression.

These data suggest that variance is a driver of stereotypical aging phenotypes at the tissue level, and that constraining specific changes caused by an increase in molecular noise during aging – such as in cell-cell communication nodes, may prevent the spread of age-related cues amongst epithelia.

### Age-dependent dysregulation in LEPs shape predictors of normal breast tissue and PAM50 subtypes

Our GSEA and literature review of genes with age-dependent changes in LEPs revealed enrichment for pathways and genes commonly dysregulated in breast cancers. Unsupervised hierarchical clustering of TCGA matched normal and primary tumor samples (n=1,201) based on the expression of these LEP-specific age-dependent DE and DV genes identified four main sample clusters (**Figure 6A**): (i) cluster 1 represented predominantly by PAM50 LumA and Her2 breast cancer subtypes; (ii) cluster 2 by PAM50 LumB and LumA subtypes; (iii) cluster 3 by PAM50 Basal subtype; and (iv) cluster 4 by matched normal samples. This suggests that age-dependent changes in LEPs may reflect dysregulation of biological processes that play a role in initiation of tumors from normal tissue, and in the etiology of breast cancer subtypes. We therefore assessed whether DE and DV genes that change in LEPs with age can be used as biomarkers that can classify normal tissue from cancer and predict breast cancer subtypes.

**Figure 6.**
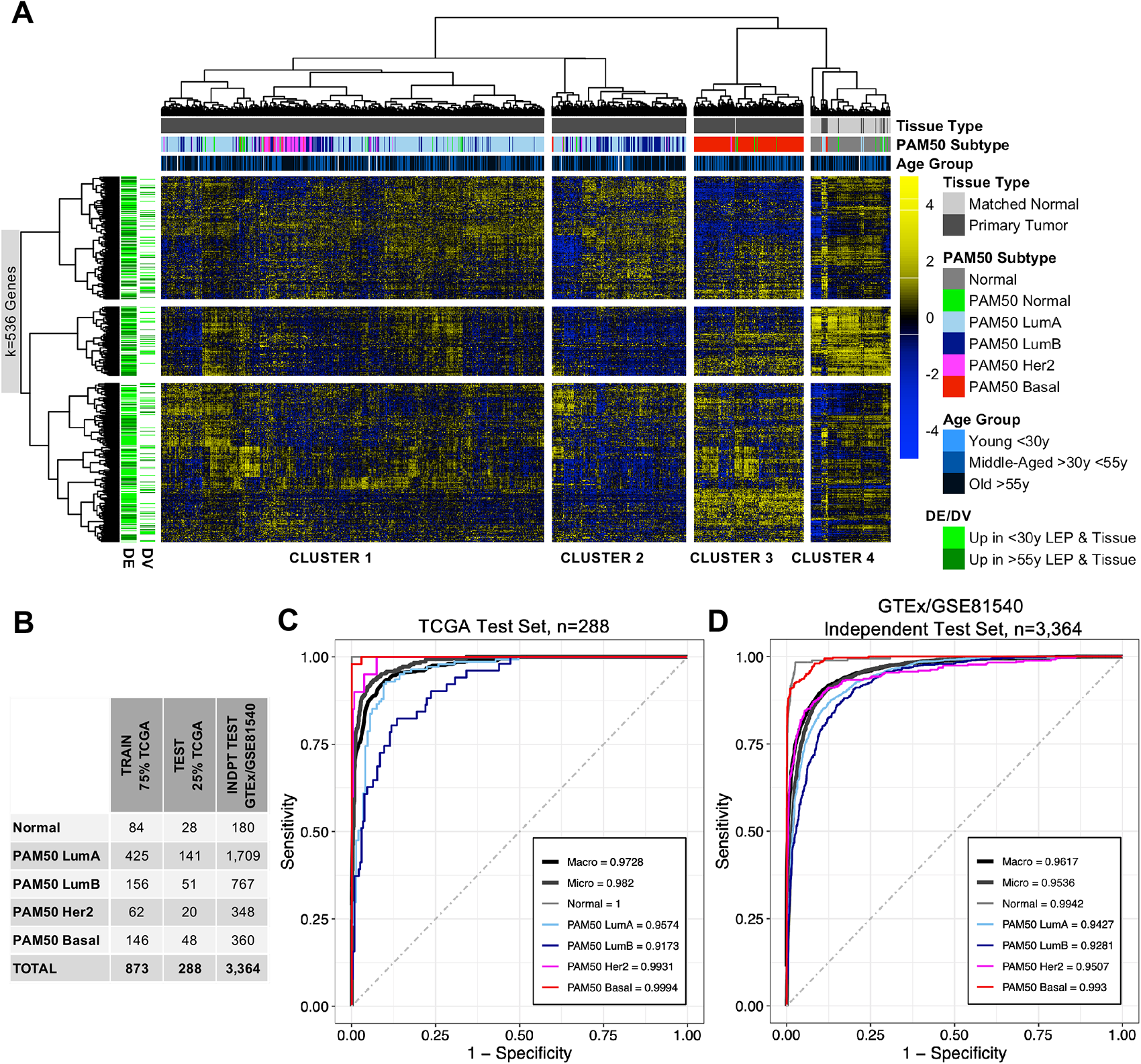
Age-dependent dysregulation in LEPs shape predictors of normal breast tissue and PAM50 subtypes. (**A**) Unsupervised hierarchical clustering of TCGA samples from matched normal and primary tumor tissue based on expression of age-dependent DE and DV genes identified in LEPs (adj. *p* < 0.05). PAM50 intrinsic subtypes and patient age at diagnosis are annotated. Gene expression scaled log_2_ FPKM values are represented in the heatmap; clustering performed using Euclidean distances and Ward agglomerative method. (Note: extreme outlier values are set to either the minimum or maximum value of the scale bar). Number of age-dependent DE and DV genes (k) included in analysis annotated. (**B**) Number of individuals in each ML class in the training, test and independent test sets. (**C-D**) Multi-class classification model performance in predicting normal tissue and breast cancer breast cancer subtypes in the (**C**) TCGA test set and (**D**) GTEx/GSE81540 independent test set. Macro AUC, micro AUC, and AUC of each group vs. rest are shown.

Using 75% of TCGA data for training and cross-validation (n=873), we built an elastic net machine learning (ML) classifier of normal breast tissue and PAM50 breast cancer subtypes based on the 536 age-dependent DE and DV genes identified in LEPs that were represented in our ML datasets. The best performing model selected during cross-validation had a mean balanced accuracy of 0.91, mean sensitivity=0.86, and mean specificity=0.96. Our ML classifier proved predictive in the remaining 25% of TCGA test data which the model had not seen (n=288, mean balanced accuracy=0.93, mean sensitivity=0.88, mean specificity=0.97), and in an independent test set composed of normal tissue from GTEx sand breast cancer tissue from GSE81540 (n=3,364, mean balanced accuracy=0.87, mean sensitivity=0.77, mean specificity=0.94) (**Figure 6B-6D**). We further assessed our ML model performance in the two test sets using three measures of the area under the curve (AUC) for multi-class prediction: (i) AUC of each group vs. the rest; (ii) micro-average AUC calculated by stacking all groups together; and (iii) macro-average AUC calculated as the average of all group results (Wei & Wang, 2020). We found all per group vs. rest AUC to be > 0.9, and micro-average and macro-average AUC > 0.95 in both the TCGA (**Figure 6C**) and GTEx/GSE81540 test sets as annotated in (**Figure 6D**). In addition to accurately classifying PAM50 subtypes, LEP-specific aging biomarkers identified normal from cancer tissue 100% and 93.3% of the time respectively in the TCGA and GTEx/GSE81540 test sets.

We next identified the genes that contributed most to the ML predictor. We identified 127 genes with scaled variable importance contribution of ≥ 25% in the prediction of at least one class (**Figure 7A**), with 18% of predictors deriving from DV analysis. Of these, estrogen receptor *ESR1* downregulated in older LEPs and transmembrane protein *TMEM45B* upregulated in older LEPs are part of the 50-gene PAM50 subtype predictors with prognostic significance (Parker et al., 2009). We highlight the top five genes with the highest variable importance for each class: (i) Normal tissue – *FN1, ABCA10, HAS3, KLHL13* and *ACVR1C*; (ii) PAM50 LumA – *FAM198B, HSD17B1,* SEC chromatin reader component *MLLT3* discussed previously*, SERPINE1* and *PLIN2*; (iii) PAM50 LumB – *GPR108, TRIM29,* desmosomal cadherin *DSG3* discussed previously*, ESR1* and *ZNRF3*; (iv) PAM50 Her2 – *ESR1, TMEM45B, FA2H, CDK12* and *TMEM63C*; and (iv) PAM50 Basal – *SLC25A37*, *PGBD5*, *BAIAP2L2*, *TBX19* and *GPR161* (**Figure 7—figure supplement 1A-1E**). Top predictors of normal tissue and of PAM50 Her2 and PAM50 Basal subtypes showed larger differences in median expression relative to other groups. In contrast PAM50 LumA and LumB top predictors showed large differences in median expression compared to non-luminal subtypes but exhibited relatively smaller but nonetheless significant differences in median expression relative to each other (**Figure 7—figure supplement 1A-1E**). While expression of these predictive genes may not be specific to the LEP lineage alone, our findings suggest that age-dependent dysregulation of these genes in LEPs could disrupt lineage-specific signaling and the homeostatic control mechanisms of key biological processes that have been implicated in breast cancers.

**Figure 7.**
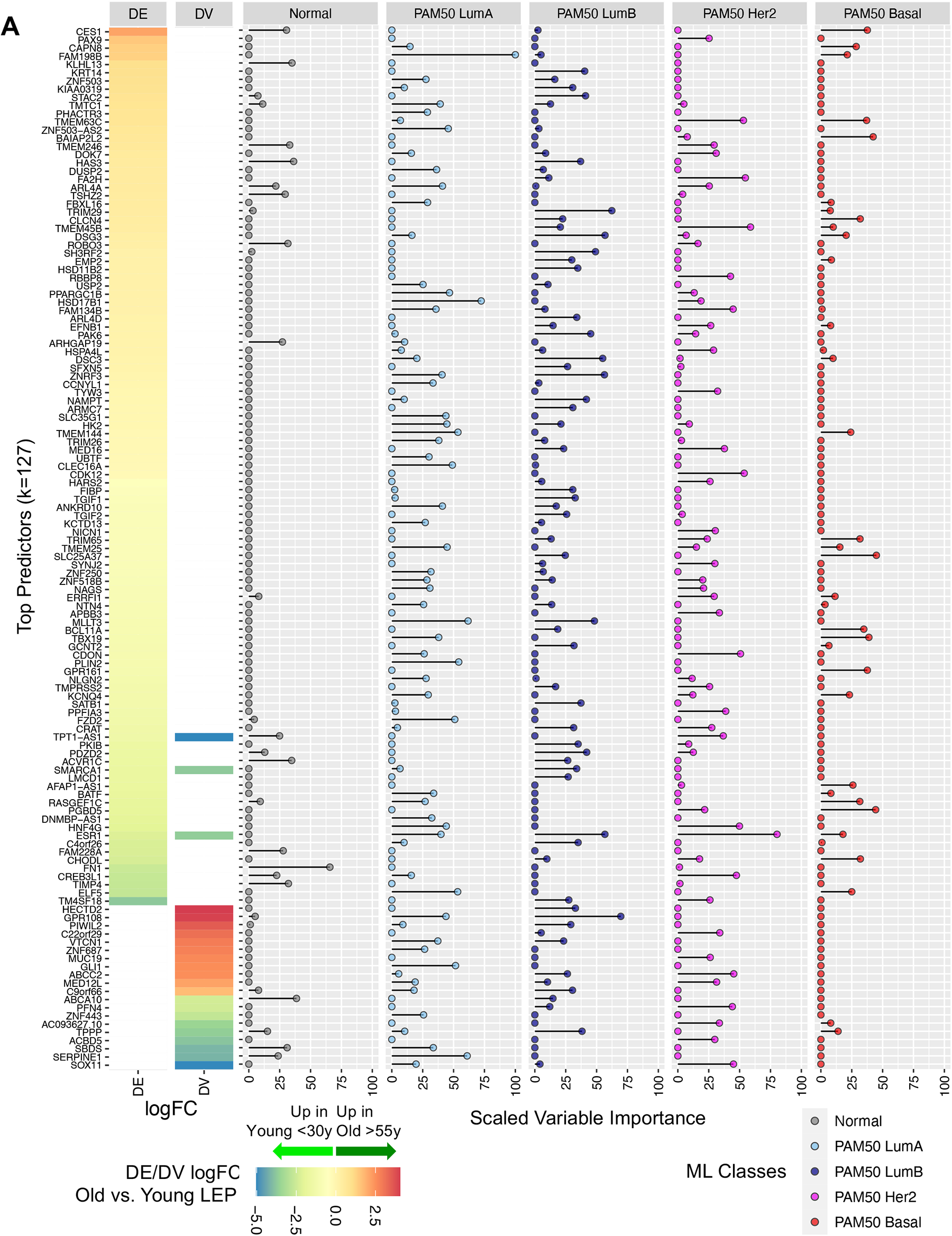
Expression of top aging-associated predictors of normal breast tissue and PAM50 subtypes in TCGA. (**A**) Gene predictors with scaled variable importance ≥ 25% in prediction of at least one class: normal breast tissue, PAM50 LumA, PAM50 LumB, PAM50 Her2 or PAM50 Basal, in TCGA. Rank ordered heatmap DE and DV lfc in LEPs with (+) lfc higher in older and (-) lfc higher in younger LEPs (left); scaled variable importance of each gene in each TCGA class (right). Number of gene predictors (k) annotated.

Our results illustrate that age-dependent changes in LEPs embody biology that is relevant and contributes to tissue-level biology predictive of breast cancer subtypes. These changes may reflect age-related dysregulation convergent with development of frank tumors. The degree and variability of these age-dependent changes across individuals may explain the differential susceptibility of specific individuals to breast cancer initiation, and the development of specific breast cancer subtypes.

## Discussion

Our analyses have shown that aging involves integration of directional and variant responses that reshape the transcriptomic landscape of the two main epithelial lineages of the breast, the LEPs and MEPs. These changes lead to a loss in lineage fidelity with age where faithfulness of lineage-specific expression is diminished. This is seen via the genome-wide loss of tuned windows of expression of lineage-specific genes and a decrease in the magnitude of differential expression between the genes that define LEPs and MEPs. Our approach delineated the contribution of each epithelial lineage and identified two models mediating this loss of lineage fidelity in breast epithelia with age – either via directional changes, as measured by differential gene expression; or via an increase in variance, as measured by differential variability analysis. Aging-dependent expression variances occur in both lineages, whereas the overwhelming majority of directional expression changes with age occurred in LEPs. This is a striking finding when one considers the two lineages arise from common progenitors, and with their juxtaposition in tissue allowing for direct cell-cell communication.

We hypothesize that LEPs are the nexus of integration for variant responses that lead to stereotyped directional age-dependent outcomes in mammary gland – the likely result of a dynamic process of iterative feedback between LEPs and MEPs, and other cell types in the breast. LEPs from older women still maintain canonical LEP-specific features, whereas they exhibit genome-wide loss of lineage fidelity that implicate dysregulation of genes with known roles in breast cancer. This suggests that susceptibility to cancer entails loss of proper specification of the luminal lineage, and that age-dependent molecular changes in LEPs contributes to this loss. Age-dependent directional changes in LEPs are detectable in bulk tissue and implicated downregulation of chromatin and genome organizers such as *SATB1*, suggesting means by which loss of lineage fidelity may be perpetuated genome-wide. The pathways affected by transcriptomic changes during aging are commonly linked with breast cancer, and the age-dependent changes in LEPs reflect relevant biology that distinguish normal tissue from breast cancers and is predictive of PAM50 breast cancer subtypes. Together our findings illustrate how age-dependent changes in LEPs contribute to the aging biology of breast tissue, and we propose that this biology reflects dysregulation convergent with processes associated with breast cancers.

Aging studies of gene expression in human tissues have been largely restricted to analysis of bulk tissue, lacked cell-type specific resolution, and were focused on directional changes with age using DE analysis. Bulk analyses make it impossible to separate impacts of aging that are driven by the intrinsic changes that occur molecularly within each lineage versus compositional changes that reflect shifts in cell type proportions. Lineage-specific analyses provide intermediate resolution between bulk RNA-seq and single-cell RNA-seq and allows for cost effective analysis of cell population-level responses and interactions. As such, lineage-resolution analyses also provide an avenue to validate computational deconvolution methods that have emerged to extract cell-type specific contributions in bulk tissue (Shen-Orr & Gaujoux, 2013; Titus et al., 2017). We provided evidence that tissue-level changes with age are driven not just by changing compositions of the breast (Benz, 2008; Garbe et al., 2012), but by intrinsic molecular changes in the underlying cell populations. Indeed, while bulk tissue expression reflects cellular heterogeneity, we were able to identify age-dependent changes in bulk tissue that mirror the DE in LEPs with age, suggesting that LEPs contribute, if not drive, the certain emergent properties of breast tissue.

In addition to down regulation of genome organizer *SATB1* in LEPs and breast tissue (adj. *p*<0.05), we also identified another 60 genes that showed concordant directional changes in LEPs (adj. *p*<0.05) and in primary tissue (unadj*. p*<0.05) despite the difference in platforms (RNA-seq vs. microarray). Network and community analyses showed enrichment of genes involved in chromosomal rearrangement, including *BCL11A –* a subunit of the BAF (SWI/SNF) chromatin remodeling complex (Kadoch et al., 2013), *MLLT3 –* a chromatin reader component of the SEC (Moustakim et al., 2018), and *PHC1* and *PCGF3* – components of the PcG multiprotein PRC-1-like complex required for developmental maintenance of repressed but transcriptionally poised chromatin configuration through alteration of chromatin accessibility, folding and global architecture of nuclear organization (Illingworth, 2019). Of note, *SATB1* was downregulated in PAM50 LumA, LumB and Her2 breast cancer subtypes relative their matched normal tissue and was expressed the lowest in luminal subtype cancers – the subtype most associated with aging. *SATB1* has genome organizing functions in tumor progression (Kohwi-Shigematsu et al., 2013), and has been described as a key regulator of EMT in cancers (Naik & Galande, 2019); however its role in normal breast epithelia remain to be elucidated. *BCL11A*, *MLLT3, ZNF521, PHC1* and *PCGF3* also show subtype-specific dysregulation in breast cancers, with *BCL11A*, *ZNF521 and PHC1* specifically downregulated in luminal subtype cancers. We speculate that chromatin and genome organization play a key role in the maintenance of the luminal lineage, and that their dysregulation may mediate loss of lineage fidelity observed genome-wide.

The striking phenotypic changes in LEPs are starkly juxtaposed to MEPs, which have so far revealed few obvious signs of changes with age. Nevertheless, heterochronous bilayers of MEPs and LEPs suggest that the chronological age of MEPs controls the biological age of LEPs, illustrating that MEPs do change with age and revealing the existence of a non-cell autonomous mechanism that integrates aging-imposed damage across the tissue (Miyano et al., 2017). Here, we showed that changes in MEPs largely involved changes in gene expression variances with age. Aging-associated increases in variances in both lineages drove a large fraction of the observed loss of lineage fidelity in epithelia of older women, comparable to the contribution of age-dependent directional changes. In our opinion, changes in variance are an underappreciated component of aging analyses. DE analysis is a ubiquitous statistical tool used in the analysis of expression profiling studies, whereas only more recently have changes in variance been systematically analyzed (Bashkeel et al., 2019; de Jong et al., 2019; Slieker et al., 2016; Xie et al., 2011), along with the development of differential variance analytical tools (Phipson et al., 2016; Phipson & Oshlack, 2014; Ran & Daye, 2017).

For changes to be detected as significant in DE analysis, the following assumptions must be met: (i) the biological phenomenon causes dysregulation that is directional – e.g., genes are either up- or down-regulated; and (ii) dysregulation occurs at the same time – e.g., in the same genes in the same pattern, across the majority of individuals in the group of interest – i.e., it is stereotypic. While some aging processes may be deterministic, like telomere shortening, other processes may be stochastic, born out of the accumulation of random physicochemical insults that manifest as an increase in noise in the system (Todhunter et al., 2018). In the latter case, the signal itself is the noise in the system. Another way to view this type of dysregulation is by observing the deviation from a set range. A change in the dynamic range of expression, for instance, of regulatory genes such as TFs that have very tuned or narrow windows of expression, can lead to dysregulation as expression deviates from the set range. This noise can lead to decoupling of tightly regulated networks. While both increase and decrease in dynamic range with age do occur, here we specifically focused on increases in variance and its effect on the loss of lineage fidelity with age, hypothesizing that genes that have very tightly tuned windows of expression in younger healthy individuals and that see large increases in variance in older subjects, are good candidates for susceptibility factors that could be predictive of breast cancer risk. Accordingly, in single-cell studies, aged cells were shown to have increased transcriptional variability and loss of transcriptional coordination compared to younger cells of the same tissue (Enge et al., 2017; Kowalczyk et al., 2015; Levy et al., 2020; Martinez-Jimenez et al., 2017), suggesting that increase in cellular heterogeneity with age underlies the population-level increases in variances between the individuals we observed. As such, the molecular signals of aging cells may not be fully captured as stereotyped directional changes – rather a large fraction of age-associated changes will be reflected as increases in measured variance in the molecular signal across an aged cohort.

We identified potential ligand-receptor pairs and junctional proteins including tight junctions, desmosomes, and gap junction components, that mediate dysregulated cell-cell and cell-microenvironment signaling within the epithelium. We provided experimental validation for the role of gap junction protein, Connexin-30 (*GJB6*) in mediating the ability of MEPs to impose an aging phenotype on LEPs (Miyano et al., 2017). It is unclear whether this occurs chemically through passage of ions or small molecules through gap junction channels, indirectly via gap junction-mediated structural proximity of LEPs and MEPs, or via signaling complexes with connexin-interacting proteins including cytoskeletal elements, tight and adherens junctions, and enzymes like kinases and phosphatases (Dbouk et al., 2009). How this occurs will require further exploration.

GSEA further identified age-dependent enrichment of gene sets in LEPs and MEPs that were commonly dysregulated in breast cancers including gene sets related to: inflammation and immunosenescence, processes synonymous with aging and cancer progression (Fulop et al., 2017); cell-cycle related targets of E2F transcription factors, which are thought to play a role in regulating cellular senescence (Lanigan et al., 2011); and targets of the oncogene *MYC.* Aging associated changes in the immune response were further implicated in our ligand-receptor pair analysis, where known immune-associated ligands and receptors exhibited loss of lineage-specific expression in breast epithelia. We showed previously that *in situ* innate and adaptive immune cell infiltration of the breast epithelia and interstitial stroma change with age (Zirbes et al., 2021) how this is linked to the dysregulation of epithelial signaling remain to be explored.

Whereas our age-specific analyses did not identify oncogenes that were DE between younger and older epithelia, gene set enrichment in LEPs and MEPs revealed a putative example of priming. We did not detect changes in Myc expression with age, but Myc targets were among the gene sets that were significantly enriched through DE or DV analysis. *Myc* is amplified or overexpressed in ∼35% of breast cancers and exerts pleiotropic effects across the genome (Xu et al., 2010). In the context of cancer progression, Myc is able to induce telomerase activity, which enables bypass of the replicative senescence barrier in mammary epithelial cells (Garbe et al., 2014). While we do not have evidence for the direct involvement of Myc in this context, we speculate that secondary events such as demethylation at Myc binding sites at target genes could explain the enrichment of Myc relevant signatures.

Using machine learning, we built a predictive elastic net model using age-dependent DE and DV genes identified in LEPs that was able to classify normal breast tissue from breast tumors and predicted breast cancer subtypes in test sets of publicly available normal and cancer tissue transcriptomes from more >3,000 women. This illustrates how tissue-level predictive biomarkers of breast cancer with subtype-specific expression relative to matched normal samples are dysregulated with age at the cell population-level as we observed in the luminal lineage. The contribution of non-epithelial cell types to the age-dependent expression of these genes in bulk tissue remain the subject of future studies. Given that the mammary epithelium is the origin of breast carcinomas and age is the most significant risk factor for breast cancers, age-dependent changes in the transcriptomic landscape of luminal cells may be a key contributing factor to the tissue-level dysregulation of cell-cell and cell-microenvironment signaling in breast cancers and may reflect relevant biology convergent with the development of frank tumors. Indeed, the variance in expression of these genes across aged individuals may reflect the differential susceptibility of certain individuals to specific breast cancer subtypes.

## Conclusions

Our studies culminate in the exploration of how directional and variant responses are integrated in breast tissue of older women that contribute to aging biology. We show that increased variance in the transcriptomic profiles of mammary epithelial lineages across individuals is a substantial outcome of aging and is likely central to our understanding of the increased susceptibility to breast cancers with age. Strikingly, LEPs are able to integrate age-dependent signals from MEPs and almost exclusively exhibit the stereotyped directional changes seen in aging epithelia that comprise a prominent signal detected in bulk tissue. Moreover, age-dependent directional and variant changes in LEPs can shape the tissue-level expression of predictive biomarkers that classify normal tissue and breast cancer subtypes, illustrating how age-dependent dysregulation may play a key role in the transition into frank cancers. We demonstrate how an increase in molecular noise during aging may lead to sufficient variance in the transcriptomes between aged individuals and propose that this is a mechanism for differential susceptibility to development of breast cancers. Because cancer susceptibility indicates a state that could be more easily pushed towards cancer initiation, we can consider the variances between aged individuals to occupy multiple metastable states, some of which represent susceptible phenotypes that can be perturbed and pushed towards cancer. We speculate that these are examples of age-dependent priming states susceptible to malignant transformation. Therefore, the degree to which breast-cancer associated genes are variably expressed across the different cell populations of the breast and across different individuals may explain why breast cancers develop in only a subset of women in a subtype-specific manner.

## List of abbreviations

18αGA: 18 alpha-glycyrrhetinic acid
adj. *p*: Adjusted *p*-value (test statistic)
AGM: Axon guidance molecule
AUC: Area under the receiving operator characteristic curve
BH: Benjamini-Hochberg
CAM: Cell adhesion molecule
ChIP: Chromatin immunoprecipitation
Cor: correlation
DE: Differential expression or differentially expressed (in context)
DV: Differential variability or differentially variable (in context)
EMT: Epithelial-to-mesenchymal transition
FDR: False discovery rate
GEO: Gene Expression Omnibus
GSEA: Gene set enrichment analysis
GTEx: The Genotype-Tissue Expression Project
HMEC: Human mammary epithelial cells
KS: Kolmogorov-Smirnov test
KW: Kruskal-Wallis test
LEP: Luminal epithelial cells
Lfc: log_2_ fold change (test statistic)
LRP: Ligand-receptor pair
MEP: Myoepithelial cells
ML: Machine learning
MSigDB: Molecular Signatures Database
PAM50: Basal PAM50 Basal-like intrinsic subtype
PAM50: PAM50 Her2-enriched intrinsic subtype
PAM50: LumA PAM50 Luminal A intrinsic subtype
PAM50: LumB PAM50 Luminal B intrinsic subtype
PAM50: Normal PAM50 Normal-like intrinsic subtype
PcG: Polycomb group
PPI: Protein-protein interaction
PRC: Polycomb repressor complex
Rlog: Regularized log
SEC: Super elongation complex
SPP: Signaling Pathways Project
STRING: Search Tool for the Retrieval of Interacting Genes/Proteins
TCGA: The Cancer Genome Atlas
TF: Transcription factor
TSS: Transcription start site

## Methods

### Experimental Model and Subject Details

The FACS-enriched luminal epithelial (LEP) and MEP myoepithelial (MEP) cells from finite lifespan, non-immortalized human mammary epithelial cells (HMECs) grown to 4^th^ passage serve as our experimental model system. HMECs were derived from normal breast tissue organoids collected from reduction mammoplasties (RM) prepared at Lawrence Berkeley National Laboratory (Berkeley, CA) with approved IRB for sample distribution and collection from specific locations. Protocols have already been established by our lab for the propagation and maintenance of these cells *in vitro* which allow for highly-reproducible source material (Garbe et al., 2009; Labarge et al., 2013).

We examined genome-wide transcription in 54 primary luminal epithelial (LEP) and MEP myoepithelial (MEP) samples from 19 women across a range of ages (**Figure 1—table supplement 1**). These epithelial lineages were isolated from human mammary epithelial cells (HMECs) from two age cohorts: younger <30y women considered to be premenopausal (age range 16-29y, m_LEP_=16, m_MEP_=16 samples, n=11 subjects) and older >55y women considered to be postmenopausal (age range 56-72y, m_LEP_=11, m_MEP_=11 samples, n=8 subjects).

### Materials Design Analysis

#### Group allocation

Samples were allocated based on age demographics of women donating tissue. As menopausal status was not available for all subjects, we used age cutoffs to identify two cohorts in this aging study. The two age groups were defined as younger <30y women considered to be premenopausal, and older >55y women considered to be postmenopausal; samples from middle-aged >30y and <55y women were excluded from analysis. Only finite lifespan cell strains derived from reduction mammoplasties from healthy women were included; strains from prophylactic mastectomies (women with high-risk breast cancer mutations or family history) or normal-adjacent to tumor tissue were excluded. No group blinding or masking was used.

#### Replicates

For the purposes of this study using finite-lifespan cell strains derived from primary tissue, subject-level biological replicates (n) refer to data derived from tissue from different subjects. Sample-level biological replicates (m) are data derived from the same subject but from separate cell culture/co-culture, FACS isolation, sample and library preparation, and RNA-sequencing experiments. Sample replicates are typically bridge samples used across different experiment batches. Technical replicates (l) are data derived from a single subject from a single sample pre-processing experiment (e.g., qPCR). When sample replication for the same subject can be modeled, analyses are done at sample level; if not, analyses are done at subject level by taking the mean value of sample replicates (see **Method Details** and **Quantification and Statistical Analysis** section). For clarity, subject (n), sample (m), and techinical (l) replicates for each analysis are annotated in each figure. The datasets generated and analyzed during the current study include RNA-sequencing count data publicly available as part of GSE182338 (Miyano et al., 2021; Sayaman et al., 2021; Shalabi et al., 2021). Criteria for inclusion of samples and gene transcripts included in the analyses, and all exclusion criteria are described in the **Method Details** section.

#### Sample-size estimation

Sample size for RNA-seq analyses was restricted to available organoids in the established HMEC bank that were isolated from reduction mammoplasties of healthy women who fall under the age range of interest: younger <30Y and older >55y, that could be expanded in primary culture. Power analyses for the limma::voom DE pipeline were conducted post-hoc using the R package ssizeRNA::check.power function (Bi & Liu, 2019) (see **Supplementary Methods**). Power calculation for lineage-specific DE between LEP and MEP with DE genes defined at BH adj*. p<*0.001 and fold-change ≥2 yielded (i) an average power (ave.pw)=0.91 and fdr average (fdr.ave)= 0.00062 in younger <30y (n=11 subjects); and (ii) ave.pw=0.90 and fdr.ave= 0.00047 in older >55y (n=8). Power calculation for age-dependent DE between young <30Y and old >55y with DE genes defined at BH adj*. p<*0.05 yielded a range of ave.pw=0.69-0.76 and fdr.ave= 0.040-0.045 in LEPs (for n=8 and n=11 respectively); (iv) ave.pw=0.87-0.89 and fdr.ave= 0.083-0.047 in MEPs (for n=8 and n=11 respectively).

#### Statistical reporting

Statistical analysis methods are described in full in the **Quantification and Statistical Analysis** section. Exact p-values are shown in figures when feasible, otherwise significance levels are annotated. Exact p-values and summary statistics at defined significance level thresholds are provided in **Supplementary Tables** (made available after publication); full summary statistics are available upon request.

### Method Details

#### Breast tissue collection and HMEC culture

Primary HMECs were initiated and maintained according to previously reported protocols using M87A medium containing cholera toxin and oxytocin at 0.5 ng/ml and 0.1nM, respectively (Garbe et al., 2009; Labarge et al., 2013). For experiments, 4^th^ passage HMECs were cultured to sub-confluence prior to FACS-sorting. HMEC strains used in this study for RNA-seq are provided (**Figure 1—table supplement 1**).

#### Flow cytometry

FACS-enriched LEPs and MEPs were isolated from 4^th^ passage finite-lifespan HMEC from reduction mammoplasties from two age cohorts: younger <30y women considered to be premenopausal (age range 16-29y) and older >55y women considered to be postmenopausal (age range 56-72y). LEP and MEP enrichment was performed across multiple studies (Miyano et al., 2021; Sayaman et al., 2021; Shalabi et al., 2021; Todhunter et al., 2021). Enrichment was conducted by FACS using well-established LEP-specific (CD227 or CD133) and MEP-specific (CD271 or CD10) cell-surface markers. Protocols were validated to sort similar populations regardless of antibody combination. Briefly, breast epithelial cells were stained and sorted following standard flow cytometry protocol. Primary HMEC strains for RNA-seq were stained with anti-human CD227-FITC (BD Biosciences, clone HMPV) or anti-human CD133-PE (BioLegend, clone7), and anti-human CD271-APC (BioLegend, clone ME20.4). Primary HMEC strains for Infinium 450K array stained with anti-human CD227-FITC (BD Biosciences, clone HMPV) and anti-human CD10-PE (BioLegnend, clone HI10a).

#### Cell co-cultures

In co-culture study (Miyano et al., 2017), FACS-enriched MEPs from 4^th^ passage HMEC were re-plated on 6-well plates and cultured until the cells were confluent. The cells were treated with Mitomycin C (Santa Cruz Biotechnology, sc-3514) at 10μg/ml for 2.5h. In co-culture with shGJB6 study, FACS-enriched control and shGJB6 transduced MEPs from older >55y women were plated on 6-well plates and cultured until the cells were confluent. FACS-enriched 4^th^ passage LEPs from younger <30y women were seeded directly on the mitomycin C-treated or shRNA transduced MEP layer. LEPs from co-cultures were separated after 10 days for gene expression qPCR analysis by FACS using anti-human CD133-PE (BioLegend, clone7) and anti-human CD271-APC (BioLegend, clone ME20.4). For Gap junction inhibition assay, cells were cultured with indicated concentration of 18-alpha-Glycryrhetinic acid (Sigma, G8503) for 7 days; LEPs from co-culture were then separated using FACS with anti-CD227-FITC (BD Biosciences, 559774, clone HMPV) and anti-CD10-PE (Biolegend, 312204, clone HI10a).

#### RNA isolation and qPCR

Total RNAs were isolated from enriched LEPs and MEPs with Quick-RNA Microprep Kit (Zymo Research, R1050). For RNA-seq, isolated RNAs were submitted to Integrative Genomic Core at City of Hope (IGC at COH) for library preparation and sequencing. For qPCR, cDNAs were synthesized with iScirpt Reverse Transcription Supermix (BioRad, 1708840) according to the manufacturer’s manual. Quantitative gene expression analysis was performed by CFX384 real-time PCR (BioRad) with iTaq Universal SYBR Green Supermix (BioRad, 1725125). Data were normalized to RPS18 or TBP by relative standard curve method.

Forward and reverse primer sequences generated in this study are indicated below: GJB6 forward and reverse primers:

5’-CTACAGGCACGAAACCACTCG-3’, 5’ACCCCTCTATCCGAACCTTCT-3’

ELF5 forward and reverse primers:

5’-TAGGGAACAAGGAATTTTTCGGG-3’, 5’-GTACACTAACCTTCGGTCAACC-3’

TBP forward and reverse primers:

5’-GAGCTGTGATGTGAAGTTTCC-3’, 5’-TCTGGGTTTGATCATTCTGTAG-3’

RPS18 forward and reverse primers:

5’-GGGCGGCGGAAAATAG-3’, 5’-CGCCCTCTTGGTGAGGT-3’

Sequences for shGJB6 and shCtrl were ggatacttgctccattcatac and gcttcgcgccgtagtctta, respectively. shCtrl (CSHCTR001LVRU6GP) and shGJB6 Lenti-virus vector (HSH06069132LVRU6GP) were purchased from GeneCopoeia.

#### RNA-sequencing

Transcriptomic profiling of LEPs and MEPs from two age cohorts: younger <30y (m=32 LEP and MEP samples, n=11 subjects) and older >55y (m=22, n=8) women (**Figure 1—table supplement 1**) was performed via RNA-sequencing as part of the LaBarge sequencing collection GSE182338 (Miyano et al., 2021; Sayaman et al., 2021; Shalabi et al., 2021). Briefly, RNA sequencing libraries were prepared with Kapa RNA mRNA HyperPrep Kit (Kapa Biosystems, Cat KR1352) or KAPA stranded mRNA-seq (Kapa Biosystems, Cat KK8420) according to the manufacturer’s protocol using 100 ng of total RNA from each sample for polyA RNA enrichment. Sequencing was performed on Illumina HiSeq 2500 with single read mode, and real-time analysis was used to process the image analysis. RNA-sequencing reads were trimmed using Trimmomatic (Bolger et al., 2014), and processed reads were mapped back to the human genome (hg19) using TOPHAT2 software (Kim et al., 2013). HTSeq (Anders & Huber, 2010) and RSeQC (Wang et al., 2012) were applied to generate the count matrices.

RNA-sequencing data pre-processing was conducted in *DESeq2* (Love et al., 2014) and *edgeR* (Robinson et al., 2010) on the entirety of the LaBarge sequencing collection GSE182338 (m=120 LEP and MEP samples, n=48 subjects) as described in (Miyano et al., 2021; Sayaman et al., 2021; Shalabi et al., 2021) including samples not included in this study. RNA-seq transcript Ensembl IDs were mapped to corresponding gene symbols, Entrez IDs and Uniprot IDs using *EnsDb.Hsapiens.v86* (*v2.99.0*) database (Rainer, 2017). We restricted analysis to the 17,328 genes with comparable dynamic ranges and consistent lineage-specific expression between primary organoid and 4^th^ passage LEPs and MEPs in both age cohorts (linear regression *R^2^*≥0.88 to 0.91, *p<*0.0001) (**Figure 1— figure supplement 1A-1D**). ComBat batch-adjusted regularized log (rlog) expression values (Johnson et al., 2007; Leek et al., 2020; Love et al., 2014) were used for visualization and downstream analysis.

#### Public Data Sets

For differential expression analysis in bulk normal primary breast tissue, GSE102088 (Song et al., 2017) microarray data (n=114) were downloaded from the Gene Expression Omnibus (GEO) database using the GEOquery (Davis & Meltzer, 2007). For machine learning, three data sets were used: (1) TCGA RNA-seq FPKM data from matched normal or PAM50 Normal, Luminal A (LumA), Luminal B (LumB), Her2 and Basal subtype breast cancer tissues (n=1,201) were downloaded using TCGAbiolinks (Colaprico et al., 2016) package; (2) GTEx RNA-seq count data from female subjects (n=180) were downloaded using the recount3 (Collado-Torres et al., 2017; Wilks et al., 2021) and FPKM transformed; and (3) GSE81540 (Brueffer et al., 2020; Brueffer et al., 2018; Dahlgren et al., 2021) RNA-seq FPKM data from PAM50 Normal, LumA, LumB, Her2 and Basal subtype breast cancer tissues (n=3,184) were downloaded from GEO.

### Quantification and Statistical Analysis

#### Differential Analyses

For differential analyses of LEP and MEP samples, a combination of lineage and age group was modeled. Differential expression (DE) was performed in limma voom (Law et al., 2014; Ritchie et al., 2015) on sample-level data from 17,328 genes with eBayes moderation and RNA-seq batch modeled as a covariate, and with adjustment for biological replicates. Differential variability (DV) was performed in MDSeq (Ran & Daye, 2017) on batch-adjusted subject-level data from 14,601 genes whose variances could be estimated after outlier removal. For lineage-specific DE analyses, contrasts between LEP and MEP in younger <30Y and in older >55y women were performed. Lineage-specific DE thresholds were set at Benjamini-Hochberg (BH) adjusted *p<*0.001 and log_2_ fold changes, lfc ≥1 in each age cohort. LEP-specific and MEP-specific genes were defined as those with lineage-specific DE in younger <30y women. For age-dependent analyses, contrasts between <30Y and >55y LEPs and <30Y and >55y MEPs were performed, and age-dependent directional or variant changes were defined at DE or DV BH adj. *p<*0.05 in each lineage.

Age-dependent DE analysis of normal primary breast tissue was performed on publicly available GSE102088 microarray data (n=114 subjects, <30y n=35, >30y<55y n=68, >55y n=11) (Song et al., 2017) in limma with eBayes moderation. Significant DE between age groups in bulk tissue were defined at BH adj. *p<*0.05 and nominal significance at unadj. *p*<0.05.

#### Gene Set Enrichment Analysis (GSEA)

Fast gene set enrichment analysis (fgsea) (Korotkevich et al., 2021) was used to identify age-dependent enrichment of Molecular Signatures Database (MSigDB) hallmark gene sets (Liberzon et al., 2015) in LEPs or MEPs using DE and DV rank-ordered test statistics. Enriched gene sets were defined as those with enrichment BH adj*. p<*0.05. For bulk tissue GSEA analysis (GSE102088, <30y n=35, >55y n=11), gene sets were constructed from age-dependent genes in LEPs: (i) 251 genes that were differentially upregulated in young <30y LEPs; and (ii) 220 genes that were that were differentially upregulated in old >55y LEPs. Age-dependent enrichment was assessed in bulk tissue using DE rank-ordered test statistics; enrichment was similarly defined at BH adj*. p<*0.05.

#### Lineage-specific Ligand-Receptor Pair Interactions and Functional Network Analysis

Ligand-receptor pairs (LRPs) (Ramilowski et al., 2015) gene symbols were mapped to Ensembl IDs using *EnsDb.Hsapiens.v86* database (Rainer, 2017). Lineage-specific LRPs were defined based on either the LEP-specific or MEP-specific (DE adj*. p<*0.001 and fold-change ≥2) expression of either the ligand and/or its cognate receptor in the younger cohort. Lineage-specific LRP interactions were considered to be lost in the older cohort when lineage-specific DE of the ligand and/or its cognate receptor was lost in the older cells (not DE at adj*. p<*0.001, lfc≥1). Functional network enrichment of LRPs in the younger cohort and LRPs lost in the older cohort were performed using the Search Tool for the Retrieval of Interacting Genes/Proteins (STRING) database (https://string-db.org/) and enriched KEGG pathways (false discovery rate, FDR *p<*0.05) were reported.

#### Age-dependent DE Protein-protein Interactions and Functional Network Analysis

Protein-protein interaction (PPI) analysis was performed using the STRING database (https://string-db.org/). All possible PPI are considered using all active interaction sources and setting minimum require interaction score to the lowest confidence threshold of 0.150. Network was visualized in igraph (Csárdi & Nepusz, 2006), and only the largest fully connected main network of genes was plotted. Community detection was performed on this main network using optimal community structure algorithm in igraph (Brandes et al., 2008). Each community was then analyzed in STRING for functional network enrichment (FDR *p<*0.05) and common functional terms were summarized and reported.

#### Unsupervised hierarchical clustering

Unsupervised hierarchical clustering in heatmaps (gplots::heatmap.2, pheatmap) (Kolde, 2019; Warnes et al., 2020) were implemented using hclust Ward’s clustering criterion (ward.D2) agglomerative method with Euclidean distances as distance metric.

#### Machine Learning

Machine learning (ML) multi-class prediction of normal breast tissue and PAM50 breast cancer subtypes was performed in caret (Kuhn, 2008) using an elastic net model (glmnet) (Friedman et al., 2010) based on tissue expression of 536 mapped age-dependent DE and DV genes identified in LEPs. ML was carried out in three large publicly available RNA-seq datasets of normal and cancer breast tissue: GTEx, TCGA and GSE81540 (Brueffer et al., 2020; Brueffer et al., 2018; Dahlgren et al., 2021) with analysis restricted to tissues from women annotated as normal or PAM50 LumA, LumB, Her2 and Basal subtypes. The ML model was trained using 10-fold cross-validation with 3 repeats in 75% of TCGA data (n=873) using a hybrid subsampling technique (SMOTE) (Torgo, 2010), and optimizing for mean balanced accuracy. Model performance was then evaluated in the 25% of TCGA (n=288) and an independent dataset of normal tissues from GTEx and breast cancer tissues from GSE81540 (n=3,364). ML multi-class prediction performance was evaluated in each test set using the MultiROC package (Wei & Wang, 2020): (i) macro-average area under the ROC curve (AUC), calculated as the average of all group results; (ii) micro-average AUC, calculated by stacking all groups together; and (iii) AUC of each group vs. the rest. Gene predictors were identified as genes with scaled variable importance contribution to the predictive model. Genes with scaled variable importance ≥25% in prediction of at least one class were visualized; gene expression the top 5 predictors in each class were further analyzed in the TCGA breast cancer cohort.

For a full description of the **Materials Design Analysis**, **Method Details**, and **Quantification and Statistical Analysis** sections, see **Supplementary Methods**.

## Declarations

### Availability of data and materials

Human mammary epithelial cells (HMECs) derived from subjects included in this study are available upon request. Forward and reverse primer sequences generated in this study are provided in the Methods section. The datasets generated and analyzed during the current study include RNA-sequencing count data publicly available as part of GSE182338 (Miyano et al., 2021; Sayaman et al., 2021; Shalabi et al., 2021). The gene expression data that support the findings of this study are available from GSE102088 (Song et al., 2017); GSE81540 (Brueffer et al., 2020; Brueffer et al., 2018; Dahlgren et al., 2021); The Cancer Genome Atlas (TCGA) Research Network: https://www.cancer.gov/tcga; and The Genotype-Tissue Expression (GTEx) Project: https://gtexportal.org/. Analysis was conducted using standard R/Bioconductor packages and statistical tests implemented in R. All package versions, model design, and parameters are described in detail in **Supplementary Methods**. Summary statistics at defined significance levels are provided in **Supplementary Tables** (made available after publication); full summary statistics are available upon request.

### Authors’ contributions

Conceptualization, RWS, MM, MAL; Methodology, RWS, MM; Software Programming, RWS; Validation, MM, RWS, PS; Formal Analysis, RWS, MM; Investigation, RWS, MM, MRS, SS, PS, MET, AZ; Resources, MAL, MRS, MM; Data Curation, MRS, MM, RWS, MET, SS; Writing – Original Draft Preparation, RWS, MAL, MM; Writing – Review & Editing Preparation, RWS, MAL, MM, DES, MRS, PS, SLN, AZ, MET, VS, SS; Visualization, RWS, MM; Supervision, MAL; Project Administration, MAL; Funding Acquisition, MAL.

## Acknowledgments

We would like to acknowledge the contributions of Dr. James Garbe to the HMEC bank; our current and previous research associates Jennifer Lopez, Jessica Bloom and Jonathan Lee for their technical support; our patient advocates, Susan Samson and Sandy Preto; and the City of Hope Bioinformatics Core led by Dr. Xiwei Wu, specifically Dr. Min-Hsuan Chen who generated the RNA-sequencing raw count data, and Dr. Jinhui Wang who prepared the RNA-sequencing library and performed the sequencing on the Illumina HiSeq2500 platform. The results

## Supplementary Files

**File name:** Sayaman_Miyano_etal_bioRxiv2022_Supplementary_Methods

**File format:** .pdf

**Title of data: Supplementary Methods**

**Description of data:** A full description of the **Method Details** and **Quantification and Statistical Analysis** section of the **Methods.**

**File name:** Sayaman_Miyano_etal_bioRxiv2022_Supplementary_Table1

**File format:** .pdf

**Title of data: Figure 1—table supplement 1. RNA-sequencing sample list.**

**Description of data:** Metadata for RNA-sequencing samples used in this study.

**Supplementary Tables** related to Figures 1-7 will be made available upon publication.

