## Supplementary Table 1 for "Luminal epithelial cells integrate variable responses to aging into stereotypical changes that underlie breast cancer susceptibility"

| RNASeqID | SubjectID | SampleID | SampleType | CultureCondition | CellType | Age | AgeGroup | TissueType | RNASeqBatch |
|---|---|---|---|---|---|---|---|---|---|
| 14897_124_LEP_RNA_CGTACG_L999.bam.count | 124 | 124 | 124 LEP | p4 | Luminal epithelial cells | 29 | Young | Reduction Mammaplasty | Miyano1 |
| 25426_RNA_124_LEP_S36_L999.bam.counts | 124 | 124 | 124 LEP | p4 | Luminal epithelial cells | 29 | Young | Reduction Mammaplasty | Shalabi1 |
| 14896_124_MEP_RNA_GTTTCG_L999.bam.count | 124 | 124 | 124 MEP | p4 | Myoepithelial cells | 29 | Young | Reduction Mammaplasty | Miyano1 |
| 25425_RNA_124_MEP_S35_L999.bam.counts | 124 | 124 | 124 MEP | p4 | Myoepithelial cells | 29 | Young | Reduction Mammaplasty | Shalabi1 |
| 14919_160_LEP_RNA_ATTCCT_L999.bam.count | 160 | 160 | 160 LEP | p4 | Luminal epithelial cells | 16 | Young | Reduction Mammaplasty | Miyano1 |
| 14918_160_MEP_RNA_ACTGAT_L999.bam.count | 160 | 160 | 160 MEP | p4 | Myoepithelial cells | 16 | Young | Reduction Mammaplasty | Miyano1 |
| 33667_163_LEP_S6_L999.bam.count | 163 | 163 | 163 LEP | p4 | Luminal epithelial cells | 27 | Young | Reduction Mammaplasty | Todhunter1 |
| 33666_163_MEP_S5_L999.bam.count | 163 | 163 | 163 MEP | p4 | Myoepithelial cells | 27 | Young | Reduction Mammaplasty | Todhunter1 |
| 27013_168R_LEP_S33_L999.bam.count | 168R | 168R | 168R LEP | p4 | Luminal epithelial cells | 19 | Young | Reduction Mammaplasty | Miyano2 |
| 27003_168R_MEP_S24_L999.bam.count | 168R | 168R | 168R MEP | p4 | Myoepithelial cells | 19 | Young | Reduction Mammaplasty | Miyano2 |
| 14909_172L_LEP_RNA_AGTTCC_L999.bam.count | 172L | 172L | 172L LEP | p4 | Luminal epithelial cells | 28 | Young | Reduction Mammaplasty | Miyano1 |
| 14908_172L_MEP_RNA_AGTCAA_L999.bam.count | 172L | 172L | 172L MEP | p4 | Myoepithelial cells | 28 | Young | Reduction Mammaplasty | Miyano1 |
| 27014_184D_LEP_S34_L999.bam.count | 184D | 184D | 184D LEP | p4 | Luminal epithelial cells | 21 | Young | Reduction Mammaplasty | Miyano2 |
| 25481_184D_LEP_S35_L999.bam.counts | 184D | 184D | 184D LEP | p4 | Luminal epithelial cells | 21 | Young | Reduction Mammaplasty | Shalabi2 |
| 27004_184D_MEP_S25_L999.bam.count | 184D | 184D | 184D MEP | p4 | Myoepithelial cells | 21 | Young | Reduction Mammaplasty | Miyano2 |
| 25480_184D_MEP_S34_L999.bam.counts | 184D | 184D | 184D MEP | p4 | Myoepithelial cells | 21 | Young | Reduction Mammaplasty | Shalabi2 |
| 14899_240L_LEP_RNA_ACTGAT_L999.bam.count | 240L | 240L | 240L LEP | p4 | Luminal epithelial cells | 19 | Young | Reduction Mammaplasty | Miyano1 |
| 27010_240L_LEP_S29_L999.bam.count | 240L | 240L | 240L LEP | p4 | Luminal epithelial cells | 19 | Young | Reduction Mammaplasty | Miyano2 |
| 25430_RNA_240L_LEP_S4_L999.bam.counts | 240L | 240L | 240L LEP | p4 | Luminal epithelial cells | 19 | Young | Reduction Mammaplasty | Shalabi2 |
| 33663_240LB_LEP_S2_L999.bam.count | 240L | 240LB | 240LB LEP | p4 | Luminal epithelial cells | 19 | Young | Reduction Mammaplasty | Todhunter1 |
| 14898_240L_MEP_RNA_GAGTGG_L999.bam.count | 240L | 240L | 240L MEP | p4 | Myoepithelial cells | 19 | Young | Reduction Mammaplasty | Miyano1 |
| 27000_240L_MEP_S10_L999.bam.count | 240L | 240L | 240L MEP | p4 | Myoepithelial cells | 19 | Young | Reduction Mammaplasty | Miyano2 |
| 25429_RNA_240L_MEP_S3_L999.bam.counts | 240L | 240L | 240L MEP | p4 | Myoepithelial cells | 19 | Young | Reduction Mammaplasty | Shalabi2 |
| 33662_240LB_MEP_S1_L999.bam.count | 240L | 240LB | 240LB MEP | p4 | Myoepithelial cells | 19 | Young | Reduction Mammaplasty | Todhunter1 |
| 27012_356E_LEP_S32_L999.bam.count | 356E | 356E | 356E LEP | p4 | Luminal epithelial cells | 21 | Young | Reduction Mammaplasty | Miyano2 |
| 27002_356E_MEP_S19_L999.bam.count | 356E | 356E | 356E MEP | p4 | Myoepithelial cells | 21 | Young | Reduction Mammaplasty | Miyano2 |
| 27011_48R_LEP_S30_L999.bam.count | 48R | 48R | 48R LEP | p4 | Luminal epithelial cells | 16 | Young | Reduction Mammaplasty | Miyano2 |
| 27001_48R_MEP_S11_L999.bam.count | 48R | 48R | 48R MEP | p4 | Myoepithelial cells | 16 | Young | Reduction Mammaplasty | Miyano2 |
| 14893_51L_LEP_RNA_GTCCGC_L999.bam.count | 51L | 51L | 51L LEP | p4 | Luminal epithelial cells | 27 | Young | Reduction Mammaplasty | Miyano1 |
| 14892_51L_MEP_RNA_CCGTCC_L999.bam.count | 51L | 51L | 51L MEP | p4 | Myoepithelial cells | 27 | Young | Reduction Mammaplasty | Miyano1 |
| 14907_59L_LEP_RNA_CTTGTA_L999.bam.count | 59L | 59L | 59L LEP | p4 | Luminal epithelial cells | 23 | Young | Reduction Mammaplasty | Miyano1 |
| 14906_59L_MEP_RNA_GGCTAC_L999.bam.count | 59L | 59L | 59L MEP | p4 | Myoepithelial cells | 23 | Young | Reduction Mammaplasty | Miyano1 |
| 27018_29_LEP_S38_L999.bam.count | 29 | 29 | 29 LEP | p4 | Luminal epithelial cells | 68 | Old | Reduction Mammaplasty | Miyano2 |
| 27008_29_MEP_S21_L999.bam.count | 29 | 29 | 29 MEP | p4 | Myoepithelial cells | 68 | Old | Reduction Mammaplasty | Miyano2 |
| 14901_237_LEP_RNA_CGATGT_L999.bam.count | 237 | 237 | 237 LEP | p4 | Luminal epithelial cells | 66 | Old | Reduction Mammaplasty | Miyano1 |
| 14900_237_MEP_RNA_ATTCCT_L999.bam.count | 237 | 237 | 237 MEP | p4 | Myoepithelial cells | 66 | Old | Reduction Mammaplasty | Miyano1 |
| 14895_112R_LEP_RNA_GTGGCC_L999.bam.count | 112R | 112R | 112R LEP | p4 | Luminal epithelial cells | 61 | Old | Reduction Mammaplasty | Miyano1 |
| 27015_112R_LEP_S35_L999.bam.count | 112R | 112R | 112R LEP | p4 | Luminal epithelial cells | 61 | Old | Reduction Mammaplasty | Miyano2 |
| 33665_112R_LEP_S4_L999.bam.count | 112R | 112R | 112R LEP | p4 | Luminal epithelial cells | 61 | Old | Reduction Mammaplasty | Todhunter1 |
| 14894_112R_MEP_RNA_GTGAAA_L999.bam.count | 112R | 112R | 112R MEP | p4 | Myoepithelial cells | 61 | Old | Reduction Mammaplasty | Miyano1 |
| 27005_112R_MEP_S26_L999.bam.count | 112R | 112R | 112R MEP | p4 | Myoepithelial cells | 61 | Old | Reduction Mammaplasty | Miyano2 |
| 33664_112R_MEP_S3_L999.bam.count | 112R | 112R | 112R MEP | p4 | Myoepithelial cells | 61 | Old | Reduction Mammaplasty | Todhunter1 |
| 27016_117R_LEP_S36_L999.bam.count | 117R | 117R | 117R LEP | p4 | Luminal epithelial cells | 56 | Old | Reduction Mammaplasty | Miyano2 |
| 25420_RNA_117R_LEP_S30.bam.count | 117R | 117R | 117R LEP | p4 | Luminal epithelial cells | 56 | Old | Reduction Mammaplasty | Shalabi1 |
| 27006_117R_MEP_S27_L999.bam.count | 117R | 117R | 117R MEP | p4 | Myoepithelial cells | 56 | Old | Reduction Mammaplasty | Miyano2 |
| 25419_RNA_117R_MEP_S29.bam.count | 117R | 117R | 117R MEP | p4 | Myoepithelial cells | 56 | Old | Reduction Mammaplasty | Shalabi1 |
| 14905_122L_LEP_RNA_GATCAG_L999.bam.count | 122L | 122L | 122L LEP | p4 | Luminal epithelial cells | 66 | Old | Reduction Mammaplasty | Miyano1 |
| 14904_122L_MEP_RNA_CAGATC_L999.bam.count | 122L | 122L | 122L MEP | p4 | Myoepithelial cells | 66 | Old | Reduction Mammaplasty | Miyano1 |
| 27017_153L_LEP_S37_L999.bam.count | 153L | 153L | 153L LEP | p4 | Luminal epithelial cells | 60 | Old | Reduction Mammaplasty | Miyano2 |
| 27007_153L_MEP_S20_L999.bam.count | 153L | 153L | 153L MEP | p4 | Myoepithelial cells | 60 | Old | Reduction Mammaplasty | Miyano2 |
| 14911_191L_LEP_RNA_CCGTCC_L999.bam.count | 191L | 191L | 191L LEP | p4 | Luminal epithelial cells | 56 | Old | Reduction Mammaplasty | Miyano1 |
| 14910_191L_MEP_RNA_ATGTCA_L999.bam.count | 191L | 191L | 191L MEP | p4 | Myoepithelial cells | 56 | Old | Reduction Mammaplasty | Miyano1 |
| 27019_429ER_LEP_S22_L999.bam.count | 429ER | 429ER | 429ER LEP | p4 | Luminal epithelial cells | 72 | Old | Reduction Mammaplasty | Miyano2 |
| 27009_429ER_MEP_S28_L999.bam.count | 429ER | 429ER | 429ER MEP | p4 | Myoepithelial cells | 72 | Old | Reduction Mammaplasty | Miyano2 |
| 33685_124_org_LEP_S26_L999.bam.count | 124 | 124 | 124 LEP | organoid | Luminal epithelial cells | 29 | Young | Reduction Mammaplasty | Todhunter1 |
| 33684_124_org_MEP_S25_L999.bam.count | 124 | 124 | 124 MEP | organoid | Myoepithelial cells | 29 | Young | Reduction Mammaplasty | Todhunter1 |
| 33689_160_org_LEP_S32_L999.bam.count | 160 | 160 | 160 LEP | organoid | Luminal epithelial cells | 16 | Young | Reduction Mammaplasty | Todhunter1 |
| 33689_160_org_LEP_S32_L999.bam.count | 160 | 160 | 160 MEP | organoid | Myoepithelial cells | 16 | Young | Reduction Mammaplasty | Todhunter1 |
| 33681_195L_org_LEP_S22_L999.bam.count | 195L | 195L | 195L LEP | organoid | Luminal epithelial cells | 24 | Young | Reduction Mammaplasty | Todhunter1 |
| 33680_195L_org_MEP_S21_L999.bam.count | 195L | 195L | 195L MEP | organoid | Myoepithelial cells | 24 | Young | Reduction Mammaplasty | Todhunter1 |
| 33683_51L_org_LEP_S24_L999.bam.count | 51L | 51L | 51L LEP | organoid | Luminal epithelial cells | 27 | Young | Reduction Mammaplasty | Todhunter1 |
| 33682_51L_org_MEP_S23_L999.bam.count | 51L | 51L | 51L MEP | organoid | Myoepithelial cells | 27 | Young | Reduction Mammaplasty | Todhunter1 |
| 33691_237_org_LEP_S33_L999.bam.count | 237 | 237 | 237 LEP | organoid | Luminal epithelial cells | 66 | Old | Reduction Mammaplasty | Todhunter1 |
| 33687_112R_org_LEP_S30_L999.bam.count | 112R | 112R | 112R LEP | organoid | Luminal epithelial cells | 61 | Old | Reduction Mammaplasty | Todhunter1 |
| 33686_112R_org_MEP_S29_L999.bam.count | 112R | 112R | 112R MEP | organoid | Myoepithelial cells | 61 | Old | Reduction Mammaplasty | Todhunter1 |
| 33693_96L_org_LEP_S34_L999.bam.count | 96L | 96L | 96L LEP | organoid | Luminal epithelial cells | 62 | Old | Reduction Mammaplasty | Todhunter1 |
