## Supplementary Figures for "Luminal epithelial cells integrate variable responses to aging into stereotypical changes that underlie breast cancer susceptibility"

[9] Lead contact

### Figure 1—figure supplement 1

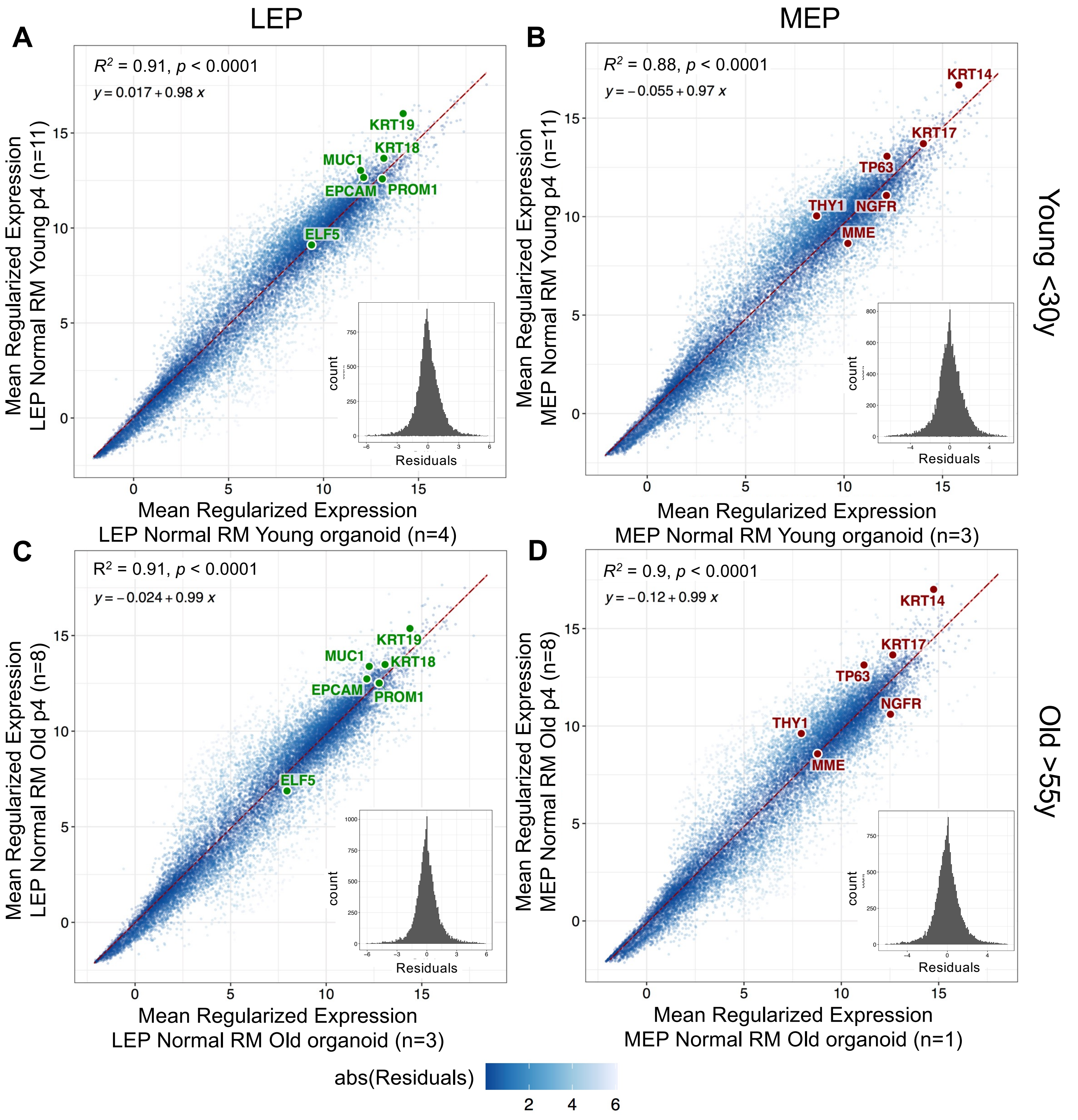


**E**

Number of Genes DE in MEP vs LEP in Young ≤ 30y

| | # Differentially Expressed | ≥ 2-fold change | ≥ 4-fold change | ≥ 8-fold change |
|---|---|---|---|---|
| adj. p-val < 0.05 | 11,265 | 4,255 | 2,025 | 1,089 |
| adj. p-val < 0.01 | 10,096 | 4,216 | 2,025 | 1,089 |
| adj. p-val < 0.001 | 8,860 | 4,040 | 2,024 | 1,089 |

**F**

Number of Genes DE in MEP vs LEP in Old ≥ 55y

| | # Differentially Expressed | ≥ 2-fold change | ≥ 4-fold change | ≥ 8-fold change |
|---|---|---|---|---|
| adj. p-val < 0.05 | 9,620 | 3,857 | 1,827 | 941 |
| adj. p-val < 0.01 | 8,028 | 3,645 | 1,824 | 941 |
| adj. p-val < 0.001 | 6,577 | 3,345 | 1,812 | 940 |

**Figure 1—figure supplement 1. Genome-wide loss of lineage-specific expression with age.** (**A-D**) Pair-wise comparison of subject-level regularized log (rlog) gene expression means between primary organoids and 4$^{th}$ passage in FACS-enriched (**A,C**) LEPs or (**B,D**) MEPs cells isolated from finite-lifespan HMECs derived from reduction mammoplasties of younger (**A,B**) or older (**C,D**) women. Linear regression line (dotted red) and standard error (red) shown with regression $R^2$, coefficient *p*-value, slope and y-intercept annotated. distribution of residuals shown in the inset.  Number of subjects (n) in each group annotated. Established lineage-specific markers are shown as reference: LEP-specific markers *KRT19, KRT18, MUC1 (CD227), EPCAM, PROM1,* and *ELF5* (green), and MEP-specific markers *KRT14, KRT17, MME (CD10), NGFR (CD271), THY1,* and *TP63* (red). (**E-F**) Table listing the number of DE genes between LEPs and MEPs in younger (**E**) and older women (**F**) women at different BH adj. *p*-value thresholds (< 0.05, 0.01, 0.001) and fold change cut-offs (≥ 2-, 4-, 8-fold change).

### Figure 2—figure supplement 1

**A**

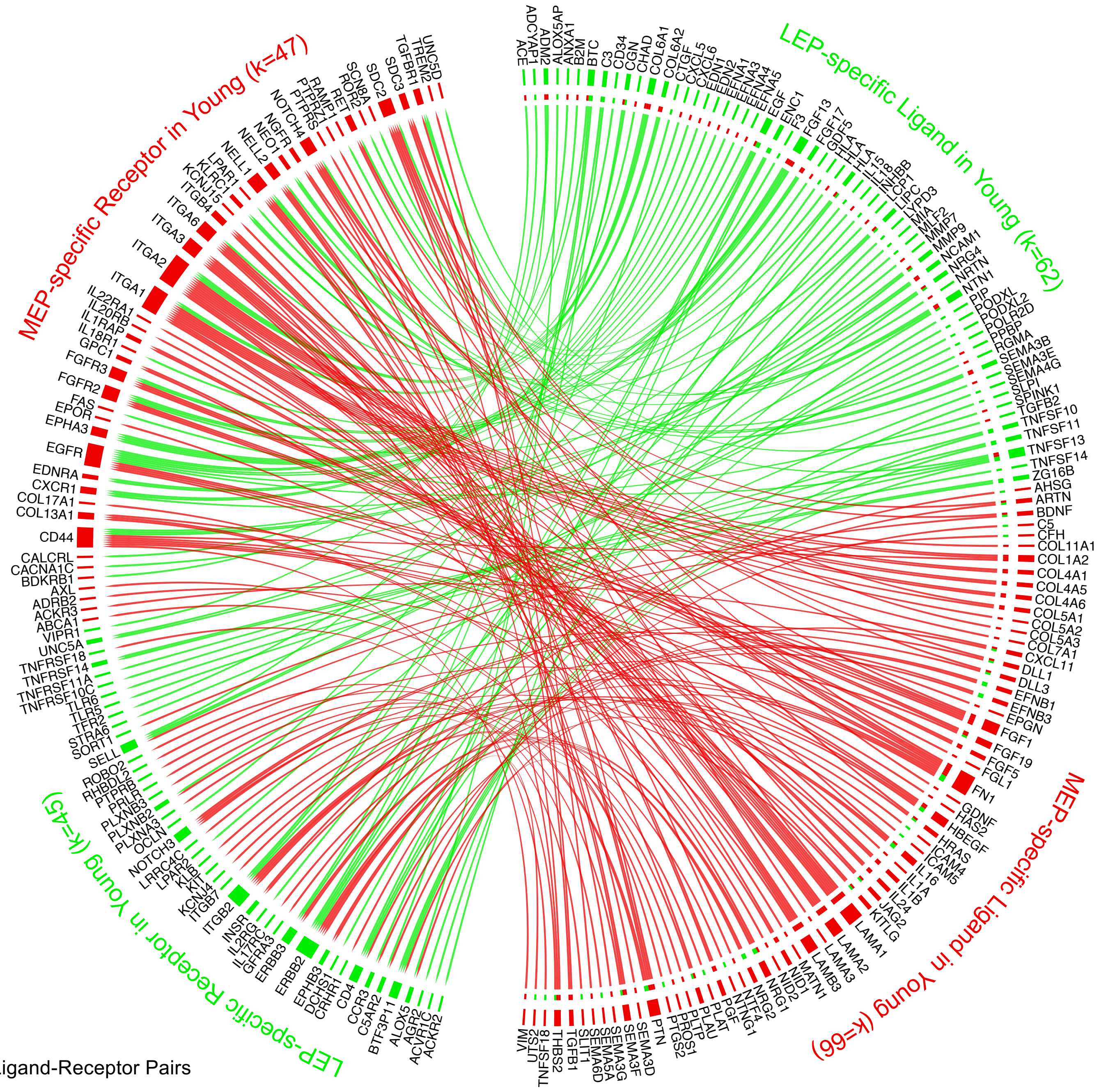


224 Ligand-Receptor Pairs

**B**

Ligand-associated KEGG Pathways

LEP

| KEGG Pathway | FDR |
|---|---|
| Cytokine-cytokine receptor interaction | 3.16E-10 |
| Rheumatoid arthritis | 8.08E-07 |
| Axon guidance | 3.66E-06 |
| PI3K-Akt signaling pathway | 4.74E-05 |
| Herpes simplex infection | 0.0006 |
| MAPK signaling pathway | 0.00081 |
| Rap1 signaling pathway | 0.00081 |
| Antigen processing and presentation | 0.00098 |
| Human papillomavirus infection | 0.00098 |
| Ras signaling pathway | 0.001 |
| Cell adhesion molecules (CAMs) | 0.001 |

MEP

| KEGG Pathway | FDR |
|---|---|
| PI3K-Akt signaling pathway | 4.02E-13 |
| Pathways in cancer | 6.93E-12 |
| Amoebiasis | 8.61E-11 |
| ECM-receptor interaction | 6.21E-10 |
| Protein digestion and absorption | 1.19E-09 |
| Focal adhesion | 2.18E-09 |
| Axon guidance | 1.04E-08 |
| Small cell lung cancer | 2.69E-08 |
| AGE-RAGE signaling pathway in diabetic complications | 3.81E-08 |
| MAPK signaling pathway | 7.23E-08 |
| Human papillomavirus infection | 1.44E-07 |
| Breast cancer | 9.83E-06 |
| Ras signaling pathway | 1.31E-05 |
| Relaxin signaling pathway | 6.39E-05 |
| Complement and coagulation cascades | 8.15E-05 |
| Endocrine resistance | 0.00019 |
| Cytokine-cytokine receptor interaction | 0.00028 |
| Toxoplasmosis | 0.00032 |
| Rap1 signaling pathway | 0.00053 |
| Leishmaniasis | 0.00079 |
| Melanoma | 0.00083 |
| Gastric cancer | 0.001 |

**C**

Receptor-associated KEGG Pathways

LEP

| KEGG Pathway | FDR |
|---|---|
| Cytokine-cytokine receptor interaction | 8.65E-07 |
| Axon guidance | 7.87E-06 |
| PI3K-Akt signaling pathway | 3.08E-05 |
| Cell adhesion molecules (CAMs) | 3.08E-05 |

MEP

| KEGG Pathway | FDR |
|---|---|
| Pathways in cancer | 1.72E-09 |
| Cytokine-cytokine receptor interaction | 4.01E-08 |
| PI3K-Akt signaling pathway | 4.01E-08 |
| Regulation of actin cytoskeleton | 5.51E-08 |
| Arrhythmogenic right ventricular cardiomyopathy (ARVC) | 7.67E-07 |
| ECM-receptor interaction | 1.24E-06 |
| Hypertrophic cardiomyopathy (HCM) | 1.24E-06 |
| Dilated cardiomyopathy (DCM) | 1.48E-06 |
| Hematopoietic cell lineage | 1.90E-06 |
| Human papillomavirus infection | 1.17E-05 |
| MAPK signaling pathway | 7.73E-05 |
| Focal adhesion | 8.48E-05 |
| Proteoglycans in cancer | 8.48E-05 |
| Central carbon metabolism in cancer | 0.0002 |
| EGFR tyrosine kinase inhibitor resistance | 0.00037 |
| Calcium signaling pathway | 0.0006 |
| Rap1 signaling pathway | 0.001 |

**Figure 2—figure supplement 1. Loss of lineage fidelity with age leads to disrupted lineage-specific signaling.** (**A**) Interactome map of the ligand-receptor pairs (LRPs) in younger women based on lineage-specific DE of ligands and their cognate receptors in LEPs (green) or MEPs (red) (adj. $p<0.001$, lfc≥1). LRPs are connected by chord diagrams from the cell type expressing the ligand (L) to the cell type expressing the cognate receptor (R). Number of LRPs, and genes (k) in each category annotated. (**B-C**) Network functional enrichment of top KEGG pathways (FDR $p<0.001$) associated with lineage-specific DE of (**B**) ligands and/or (**C**) cognate receptors in LEPs and MEPs in younger women.

### Figure 3—figure supplement 1

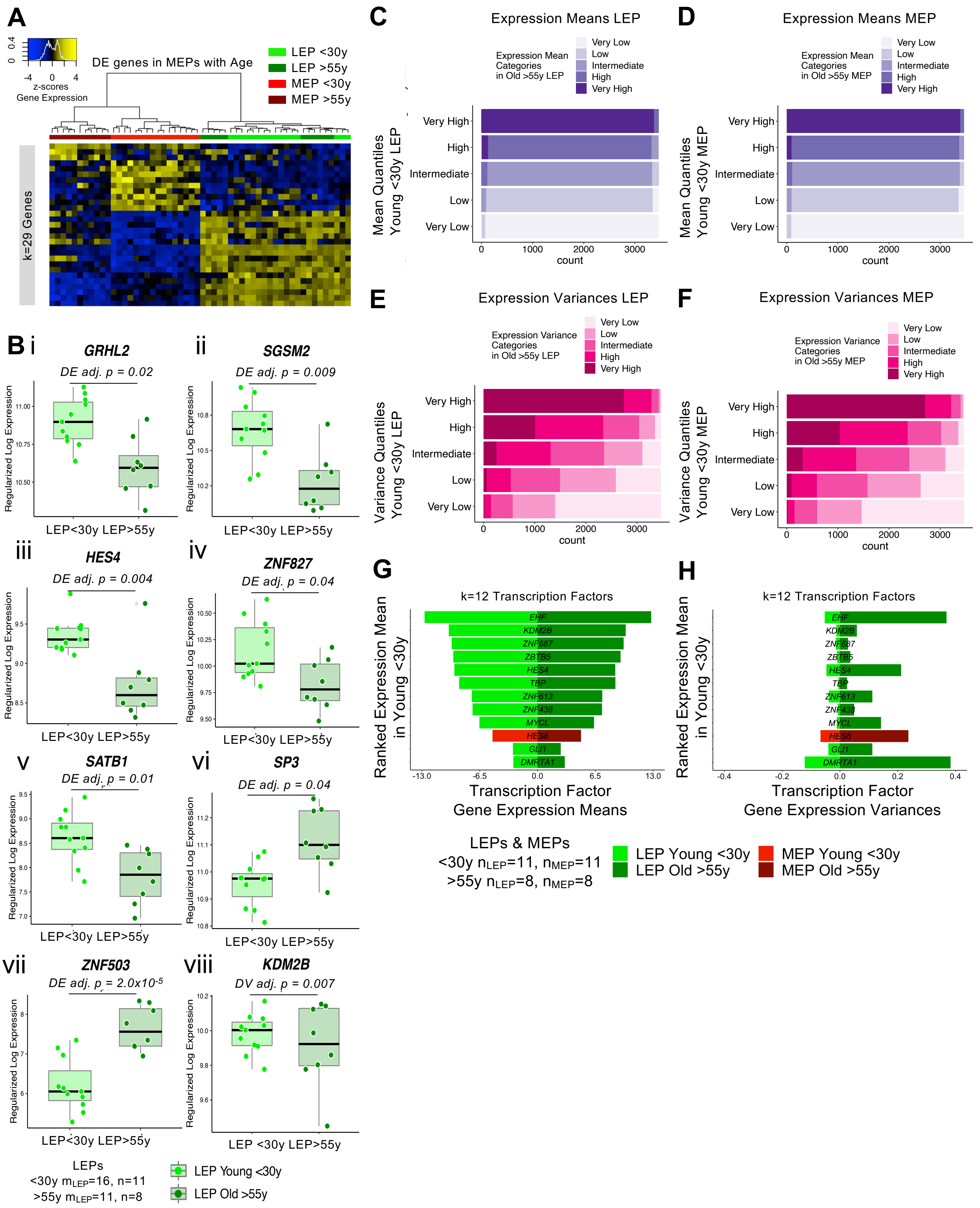

**Figure 3—figure supplement 1. The luminal lineage is a hotspot for age-dependent directional changes.** (**A**) Hierarchical clustering of all LEP and MEP samples based on sample-level expression of age-dependent DE genes in MEPs (adj. $p < 0.05$). Number of DE genes (k) indicated. Gene expression scaled regularized log (rlog) values are represented in the heatmap; clustering performed using Euclidean distances and Ward agglomerative method. (**B**) Boxplots of subject-level gene expression rlog values in younger and older LEPs of TFs: (i) *GRHL2*, (ii) *SGSM2*, (iii) *HES4*, (iv) *ZNF827*, (v) *SATB1*, (vi) *SP3* and (vii) *ZNF503* with DE adj. *p*-values; and (viii) *KDM2B* with DV adj. p-values. Number of subjects (n) and sample replicates (m) in DE analysis annotated. (**C-F**) Gene expression (**C-D**) means and (**E-F**) variances of LEPs and MEPs from younger women are categorized into quantile levels: very low, low, intermediate, high, very high. Corresponding categories of gene expression means and variances in older cells, as defined by threshold values in younger cells, are fractionally represented. (**G-H**) TF with significant increase in variances in older cells (DV adj. $p<0.05$) are shown with subject-level rlog gene expression (**G**) means in rank order from lowest to highest expressed in the younger cohort and (**H**) corresponding variances. Number of subjects (n) in analysis annotated.

### Figure 4—figure supplement 1

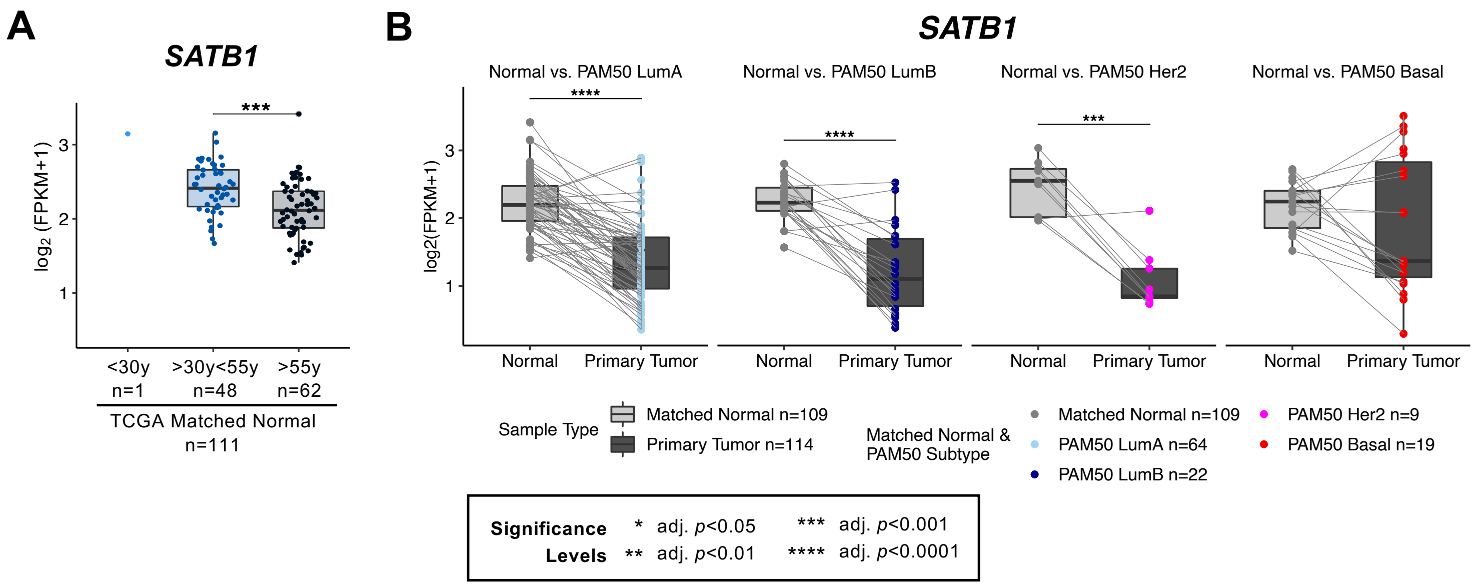

**Figure 4—figure supplement 1. Age-dependent directional changes in the luminal lineage are indicators of aging breast tissue.** (**A**) Boxplot of *SATB1* gene expression: $\log_2$ FPKM values in the TCGA breast cancer cohort by age at diagnosis in matched normal samples. Wilcoxon *p*-values between groups annotated. (**B**) Boxplots of $\log_2$ FPKM values of *SATB1* in paired matched normal and primary tumor across each PAM50 subtype in TCGA. Wilcoxon test adj. *p*-value significance annotated. Number of subjects (n) in each analysis annotated.

### Figure 4—figure supplement 2

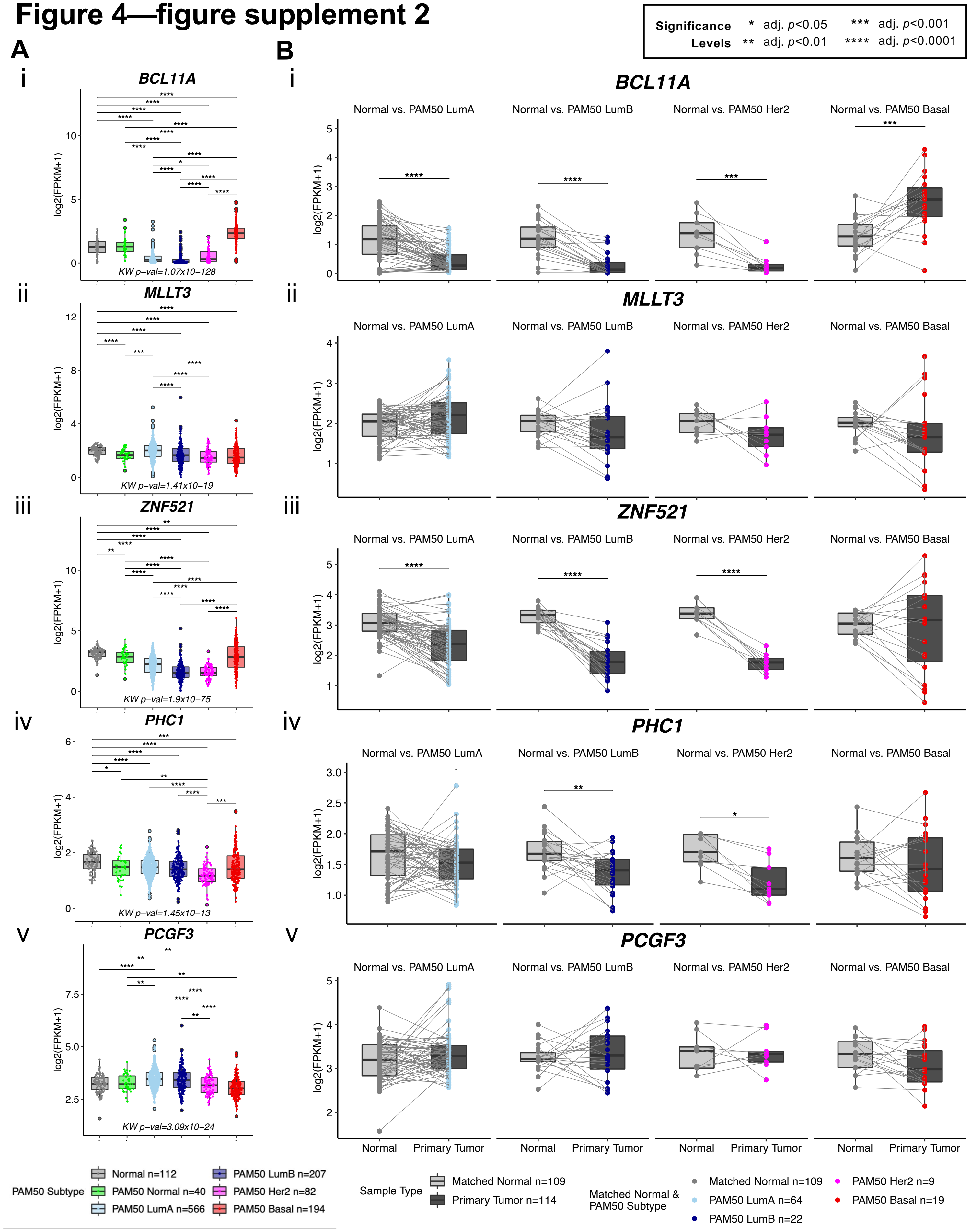

**Figure 4—figure supplement 2. Age-dependent directional changes in the luminal lineage are indicators of aging breast tissue.**

(**A**) Boxplots of $\log_2$ FPKM values of (i) *BCL11A*, (ii) *MLLT3*, (iii) *ZNF521*, (iv) *PHC1* and (v) *PCGF3* in the TCGA breast cancer cohort. Kruskal-Wallis (KW) test *p*-value between matched normal and breast cancer subtypes in TCGA; post-hoc pair-wise Wilcoxon test adj. *p*-value significance annotated. (**B**) Boxplots of $\log_2$ FPKM values of (i) *BCL11A*, (ii) *MLLT3*, (iii) *ZNF521*, (iv) *PHC1* and (v) *PCGF3* in paired matched normal and primary tumor across each PAM50 subtype in TCGA. Wilcoxon test adj. *p*-value significance annotated. Number of subjects (n) in each analysis annotated.

### Figure 5—figure supplement 1

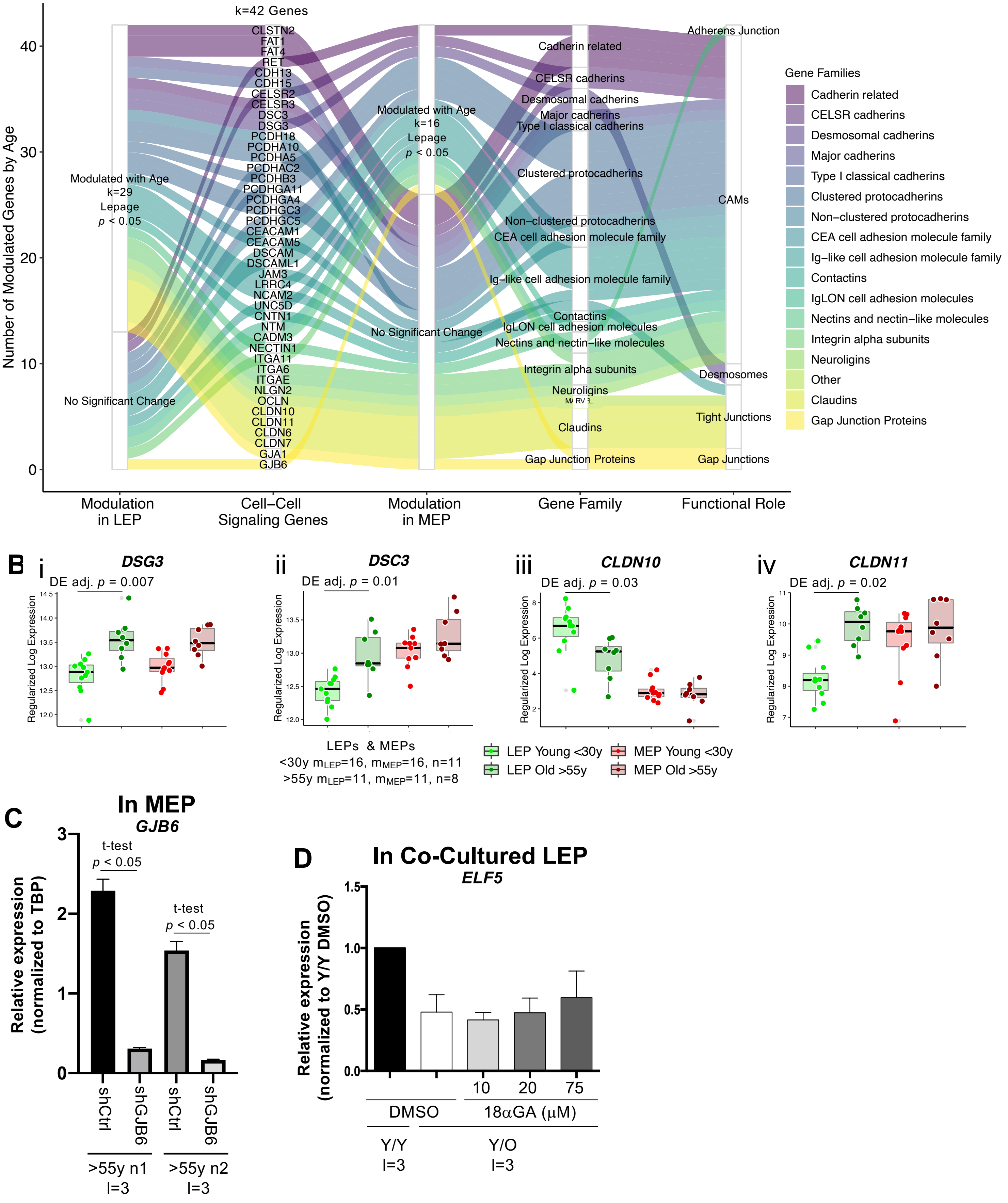

**Figure 5—figure supplement 1. Gap Junction protein *GJB6* is a mediator of the non-cell autonomous mechanism of aging in breast.** (**A**) Age-dependent modulation of apical junction-associated genes in LEPs and MEPs (Lepage test $p<0.05$). Number of age-modulated genes (k) indicated. Genes annotated with their respective HUGO Gene Nomenclature Committee (HGNC) gene family and functional role in adherens junctions, cell adhesion molecules, desmosomes, tight junctions or gap junctions. (**B**) Boxplots of subject-level rlog expression values of (i) *DSC3,* (ii) *DSG3* (iii) *CLND10*, and (iv) *CLDN11*, in LEPs and MEPs from younger and older women. Age-dependent DE adj. *p*-values are indicated. Number of subjects (n) and sample replicates (m) in DE analysis annotated. (**C**) Relative expression of *GJB6* in either shControl or shGJB6 older MEPs in two individuals. Two-tailed t-test p-value indicated. Number of subjects (n) and technical replicates (l) annotated. (**D**) Relative expression of *ELF5* in younger LEPs co-cultured with older MEPs treated with increasing concentrations of 18aGA compared to Y/Y and Y/O treated with DMSO. Number of technical replicates (l) annotated.

### Figure 7—figure supplement 1

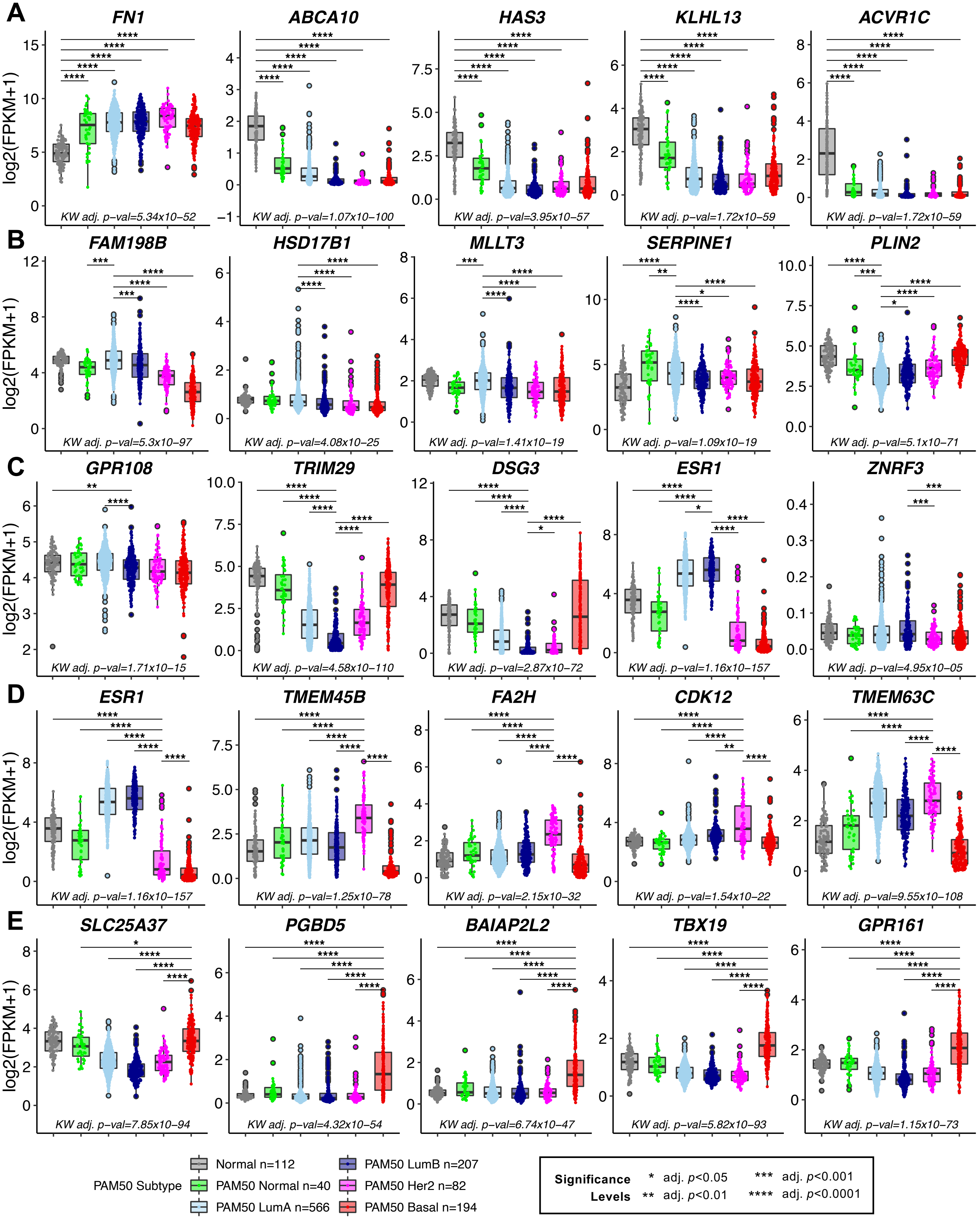

**Figure 7—figure supplement 1. Genes that distinguish normal breast tissue and PAM50 cancer subtypes show age-dependent dysregulation in luminal epithelia.** (**A-E**) Boxplots of $\log_2$ FPKM values in TCGA matched normal and PAM50 subtypes for the top five genes with the highest scaled variable importance contribution in each class in the predictive ML model: (**A**) normal tissue, (**B**) PAM50 LumA, (**C**) PAM50 LumB, (**D**) PAM50 Her2 and (**E**) PAM50 Basal. Kruskal-Wallis (KW) test *p*-value between matched normal and breast cancer subtypes in TCGA; post-hoc pair-wise Wilcoxon test adj. *p*-value significance annotated (*<0.05, **<0.01, ***<0.001, ****<0.0001). Correction for multiple hypothesis testing was performed across the five gene predictors for each class and across all pairwise comparisons; significance level shown only for pairwise comparisons to the reference class. (Note: 3 extreme outlier points in *ZNRF3* outside the plotted range in (**C**) was excluded from visualization only). Number of subjects (n) in each analysis annotated.
