## Supplementary Methods for "Luminal epithelial cells integrate variable responses to aging into stereotypical changes that underlie breast cancer susceptibility"

[9] Lead contact

**RESOURCE AVAILABILITY**

*Sample-size estimation:* Sample size for RNA-seq analyses was restricted to available organoids in the established HMEC bank that were isolated from reduction mammoplasties of healthy women who fall under the age range of interest: younger <30y and older >55y, that could be expanded in primary culture. Power analyses for the limma::voom DE pipeline were conducted post-hoc using the R package ssizeRNA (*v.1.3.2*)::check.power function which calculates average power (ave.pw) and true average FDR (ave.fdr) for a given sample size when FDR is controlled for using the BH step-up procedure (Bi & Liu, 2019). These values are calculated for four DE analyses groups: lineage-specific DE between LEP and MEP in (i) young <30y group and (ii) old >55y; and age-dependent DE between young <30y and old >55y (iii) LEPs and (iv) MEPs. Input values include: total number of genes (nGenes)=17,328; calculated means of sample-level count data in control group, with LEPs serving as control group in lineage-specific analysis and young <30y as control group in age-dependent analysis; and dispersion measures calculated from sample-level normalized values in *edgeR* (*v.3.22.5*) (Robinson et al., 2010) using the estimateCommonDisp function. Power calculation is run using 10 simulations with replacement with the following DE analysis-specific values: fdr rate of 0.001 for lineage-specific and 0.05 for age-dependent analysis; proportion of non-differentially expressed genes (nGenes – DE nGenes)/nGenes) where DE nGenes is the number of genes at defined thresholds of BH adj. $p<0.001$ and fold-change ≥2 for lineage-specific DE genes and BH adj. $p<0.05$ for age-dependent DE genes; proportion of up-regulated genes among all DE genes; fold-change values of DE genes; and sample size of treatment group where we conservatively use subject-level (n) replication size.

Power calculation for lineage-specific DE between LEP and MEP with DE genes defined at BH adj. $p<0.001$ and fold-change ≥2 yielded (i) an average power (ave.pw)=0.91 and fdr average (fdr.ave)= 0.00062 in younger <30y (n=11 subjects); and (ii) ave.pw=0.90 and fdr.ave= 0.00047 in older >55y (n=8). Power calculation for age-dependent DE between young <30y and old >55y with DE genes defined at BH adj. $p<0.05$ yielded a range of (iii) ave.pw=0.69-0.76 and fdr.ave= 0.040-0.045 in LEPs (for n=8 and n=11 respectively); (iv) ave.pw=0.87-0.89 and fdr.ave= 0.083-0.047 in MEPs (for n=8 and n=11 respectively).

*Statistical reporting:* Statistical analysis methods are described in full in the **Quantification and Statistical Analysis** section. Exact p-values are shown in figures when feasible, otherwise significance levels are annotated. Exact p-values and summary statistics at defined significance level thresholds are provided in **Supplementary Tables** (made available after publication); full summary statistics are available upon request.

### METHOD DETAILS

All computational analyses were conducted using using R (3.5.0) (R Core Team, 2018) (https://www.R-project.org/) and Bioconductor (3.7) (Huber et al., 2015) (https://www.bioconductor.org/) unless otherwise noted.

*Breast tissue collection and HMEC culture:* Primary HMECs were established and maintained according to previously reported protocol using M87A medium containing cholera toxin and oxytocin at 0.5 ng/ml and 0.1nM, respectively (Garbe et al., 2009; Labarge et al., 2013). For experiments, $4^{th}$ passage HMECs were cultured for 4-6 days (depending on strain) to sub-confluence prior to FACS-sorting. Cell cultures were fed every 2 days up to and including the day before FACS-sorting. HMEC strains used in this study were listed in (**Figure 1—table supplement 1**) for RNA-seq.

*Disassociation of uncultured cells from organoids:* For dissociation of uncultured cells from organoids, organoids were digested with 0.5% trypsin/EDTA for 10min at 37C with agitation. After trypsin treatment, organoids were disrupted by vigorous shaking for 30sec. Cells were then passed through a 40um cell strainer (BD Falcon).

*Flow cytometry:* LEP and MEP enrichment was performed across multiple studies (M. Miyano, S. Shalabi, M.E. Todhunter). Enrichment was conducted by FACS (BD FACSVantage SE,

FACSAriaIII, FACS AriaSORP or Bio-Rad S3 Cell Sorter) using well-established LEP-specific (CD227 or CD133) and MEP-specific (CD271 or CD10) cell-surface markers. Protocols were validated to sort similar populations regardless of antibody combination and enrichment methodology. Briefly, breast epithelial cells were stained and sorted following standard flow cytometry protocol. Primary HMEC strains for RNA-seq were stained with anti-human CD227-FITC (BD Biosciences, clone HMPV) or anti-human CD133-PE (BioLegend, clone7), and anti-human CD271-APC (BioLegend, clone ME20.4). Primary HMEC strains for Infinium 450K array stained with anti-human CD227-FITC (BD Biosciences, clone HMPV) and anti-human CD10-PE (BioLegnend, clone HI10a).

*Cell co-cultures:* In co-culture study (Miyano et al., 2017), FACS-enriched MEPs from 4$^{th}$ passage HMEC were re-plated on 6-well plates and cultured until the cells were confluent. The cells were treated with Mitomycin C (Santa Cruz Biotechnology, sc-3514) at 10μg/ml for 2.5h. In co-culture with shGJB6 study, FACS-enriched control shGJB6 transduced MEPs from older >55y women were plated on 6-well plates and cultured until the cells were confluent. FACS-enriched 4$^{th}$ passage LEPs from younger <30y women were seeded directly on the mitomycin C-treated or shRNA transduced MEP layer. LEPs from co-cultures were separated by FACS after 10 days for gene expression qPCR analysis. Co-cultured LEPs were stained with anti-human CD133-PE (BioLegend, clone7) and anti-human CD271-APC (BioLegend, clone ME20.4) and were separated using BD FACS ARIAIII.

For Gap junction inhibition assay, cells were cultured with indicated concentration of 18-alpha-Glycryrhetinic acid (Sigma, G8503) for 7days. LEP from co-culture was separated using BD FACSVantage with anti-CD227-FITC (BD Biosciences, 559774, clone HMPV) and anti-CD10-PE (Biolegend, 312204, clone HI10a).

*RNA isolation and qPCR:* Total RNAs were isolated from enriched LEPs and MEPs with Quick-RNA Microprep Kit (Zymo Research, R1050). For RNA-seq, isolated RNAs were submitted to Integrative Genomic Core at City of Hope (IGC at COH) for library preparation and sequencing. For qPCR, cDNAs were synthesized with iScirpt Reverse Transcription Supermix (BioRad, 1708840) according to the manufacturer's manual. Quantitative gene expression analysis was performed by CFX384 real-time PCR (BioRad) with iTaq Universal SYBR Green Supermix (BioRad, 1725125). Data were normalized to RPS18 or TBP by relative standard curve method.

*RNA-seq sequencing library preparation and sequencing with Illumina Hiseq2500:* RNA sequencing libraries were prepared with Kapa RNA mRNA HyperPrep Kit (Kapa Biosystems, Cat KR1352) or KAPA stranded mRNA-seq (Kapa Biosystems, Cat KK8420) according to the manufacturer's protocol. Briefly, 100 ng of total RNA from each sample was used for polyA RNA enrichment. The enriched mRNA underwent fragmentation and first strand cDNA synthesis. The combined $2^{nd}$ cDNA synthesis with dUTP and A-tailing reaction generated the resulting ds cDNA with dAMP to the 3' ends. The barcoded adaptors were ligated to the ds cDNA fragments. A 10-cycle of PCR was performed to produce the final sequencing library. The libraries were validated with the Agilent Bioanalyzer DNA High Sensitivity Kit and quantified with Qubit. Sequencing was performed on Illumina HiSeq 2500 with the single read mode of 51cycle. Real-time analysis (*RTA v2.2.38*) software was used to process the image analysis.

*Sequence alignment and gene counts:* RNA-Seq reads were trimmed to remove sequencing adapters using *Trimmomatic* (Bolger et al., 2014). The processed reads were mapped back to the human genome (hg19) using *TOPHAT2* software (Kim et al., 2013). *HTSeq* (Anders & Huber, 2010) and RSeQC (Wang et al., 2012) software were applied to generate the count matrices and strand information, respectively with default parameters.

*RNA-seq Pre-processing:* RNA-sequencing data pre-processing was conducted on the entirety of the LaBarge sequencing collection GSE182338 (Miyano et al., 2021; Sayaman et al., 2021; Shalabi et al., 2021) across organoids and 4$^{th}$ passage HMECs (m=120 LEP and MEP samples, n=48 subjects) including samples not included in this study. The experimental design group was defined by the combination of the culture condition (organoid, 4$^{th}$ passage), cell type (LEP, MEP) and age/risk status (normal risk RM younger <30y, normal risk RM older >55y, and PM/CLTT/PTT without or with germline mutation) of the samples. Raw counts for 34,623 transcripts from RNA-sequencing of FACS-sorted LEPs and MEPs were normalized and regularized log (rlog) transformed in *DESeq2* (*v1.20.0*) (Love et al., 2014) after removal of 30,196 transcripts with zero values across all samples. For initial QA, transformation was run blind to the design matrix (blind=TRUE). Sample QA revealed one outlier sample (subject ID 160, MEP from organoid) that was removed. Rlog transformation was repeated after outlier sample removal without blinding the transformation to the design matrix (~RNA-seq Batch + Design Group) (blind=FALSE). Rlog values were batch-adjusted using *ComBat* (*sva v.3.35.2*) (Johnson et al., 2007; Leek et al., 2020) function with the experimental design group as covariate in the model matrix (~Design Group). *ComBat* batch-adjusted rlog values were used for quality control and assessment of transcripts and filtered as discussed below.

*RNA-seq Filtering:* During quality control assessment, 5 entries (no feature, ambiguous, too low quality, not aligned, aligned not unique) were filtered out. Count data were imported into *edgeR* (*v.3.22.5*) (Robinson et al., 2010) as a *DGEList* object. Transcripts with low counts were determined using the *edgeR::filterbyExpr* function with experimental design group and batch as covariates in the design matrix (~0 + Design Group + RNA-seq Batch); 14,647 transcripts with low counts were removed. Next, genes with highly discordant expression levels between organoids and 4$^{th}$ passage LEPs and MEPs grown in 2D culture were assessed. Mean subject-level batch-adjusted rlog values were calculated for each lineage and age group. Linear regression was performed on the gene expression value means from cells isolated from organoid vs. 4th passage culture in each of the RM LEP <30y, LEP >55y, MEP <30y, MEP >55y subsets, and transcripts with absolute value of model residuals ≥6 (~4*sd) in either of the 4 subsets were considered outliers; 423 transcripts were flagged for exclusion in the DGEList object. Finally, we considered genes that did not maintain consistent lineage-specific expression between organoids and 4th passage cells grown in 2D culture. Normalization factors were calculated for the filtered data using *edgeR::calcNormFactors* function using TMM method. DE analysis in *limma voom* (*v.3.36.5*) (Law et al., 2014; Ritchie et al., 2015) was performed between LEPs and MEPs from younger <30 women in both organoids and 4th passage cells (see Quantification and Statistical Analysis). To more fully capture deviations, we used a less stringent cut-off and genes with at least lineage-specific DE significance of adj. $p$<0.1 between LEPs and MEPs in younger <30 women in either organoid and 4th passage 2D culture were identified. These 2,220 genes that did not maintain consistent lineage-specific expression between organoids and 4th passage cells in 2D culture were subsequently excluded. A final set of 17,328 genes were used for all downstream analyses of 4th passage and organoid data.

*Batch-adjustment of RNA-seq Data:* RNA-seq count data were pooled from five experiments and three studies (M. Miyano, S. Shalabi, M.E. Todhunter) conducted at different times. Each experiment was defined to be an RNA-seq batch. Batch effects were found during QC in hierarchical clustering (*hclust*) and principal component analysis (PCA) (*prcomp*) of LEP and MEP samples based on normalized rlog expression values.

For visualization and down-stream analysis of normalized expression data, rlog values were corrected for batch effects using *ComBat* – an empirical Bayes approach for adjusting data for batch effects that is robust to outliers in small sample sizes (Johnson et al., 2007). The experimental design group was defined by the combination of culture condition (organoid, 4th passage), cell type (LEP, MEP) and age group/risk group (average risk RM younger <30y, average risk RM older >55y, or higher risk PM/CLTT/PTT with or without germline mutation) and was included in the model matrix (~Design Group). *ComBat* batch-adjustment was applied in the *sva* (*v3.35.2*) package (Leek et al., 2020). *ComBat* batch-adjusted data were then filtered to remove the set of genes described above. Filtered, batch-adjusted data were checked with PCA, and linear regression analysis to confirm removal of PC association with RNA-seq batch. Hierarchical clustering was also used pre- and post-*ComBat* treatment for visualization of batch effects and the clustering of bridge samples.

Filtered *ComBat* batch-adjusted rlog values were subsetted for samples of interest and used for visualization (*ggplot v.2_3.3.3, gplots v.3.0.3::heatmap.2*) (Warnes et al., 2020; Wickham, 2016) and downstream analysis of expression values. Subject-level data were calculated as the mean value of the batch-adjusted rlog values for subjects with multiple samples.

For differential expression (DE) analysis of RNA-seq count data in *limma voom* (*v3.36.5*) (Law et al., 2014; Ritchie et al., 2015), RNA-seq batch was included in the linear model along with the above design group (~0 + Design Group + Batch). For differential variability (DV) analysis of RNA-seq count data in *MDSeq* (*v.1.0.5*) (Ran & Daye, 2017), since we were not able to model multiple samples from the same subject, *ComBatSeq* batch-adjustment (*sva_devel*) (Zhang et al., 2020) of the count data was first performed using the above design group in the model matrix. *ComBatSeq* batch-adjusted count data were then normalized using TMM method (*MDSeq::normalize.counts*). Subject-level data were generated as the mean value of *ComBatSeq* batch-adjusted normalized data for subjects with multiple samples and used in the DV analysis.

*Annotation of RNA-seq Data:* RNA-seq transcript Ensembl IDs were mapped to corresponding gene symbols, Entrez IDs and Uniprot IDs using *EnsDb.Hsapiens.v86* (*v2.99.0*) database (Rainer, 2017).

*RNA-seq Data Used in this Manuscript:* Transcriptomes in LEPs and MEPs from reduction mammoplasty HMECs at 4$^{th}$ passage (n=11 <30y, n=8 >55y) and uncultured organoids (n=4 <30y, n=3 >55y LEPs; n=3 <30y, n=1 >55y MEPs) were extracted from annotated pre-processed RNA-seq counts and rlog values from GSE182338 (Miyano et al., 2021; Sayaman et al., 2021; Shalabi et al., 2021) and used during QC analysis to determine exclusion of genes that did not maintain concordant lineage-specific expression between organoid and 4$^{th}$ passage (see *RNA-seq Filtering*). Gene expression data isolated 4$^{th}$ passage LEPs and MEPs from two age cohorts: younger <30y women considered to be premenopausal (age range 16-29y, $m_{LEP}$=16, $m_{MEP}$=16 samples, n=11 subjects) and older >55y women considered to be postmenopausal (age range 56-72y, $m_{LEP}$=11, $m_{MEP}$=11 samples, n=8 subjects) were used for downstream analysis.

*GSE102088 Data:* Normalized microarray $\log_2$ expression data, pheno data and feature data from normal primary breast tissues from n=114 women (GSE102088) (Song et al., 2017) were downloaded from the Gene Expression Omnibus (GEO) database (https://www.ncbi.nlm.nih.gov/geo/) using the *GEOquery* (*v.2.48.0*) package (Davis & Meltzer, 2007). The experimental design group was defined by age groups (younger <30y, middle aged >30y <55y, and older >55y) for differential expression analysis and visualization.

*GSE81540 Data:* Normalized expression from breast cancers with matched pheno and clinical PAM50 annotation (GSE81540) (Brueffer et al., 2020; Brueffer et al., 2018; Dahlgren et al., 2021) were downloaded from GEO. Available pre-processed data represent summed transcript FPKM values for each gene that were log2 transformed (FPKM+0.1). Raw data were not provided. For Machine Learning samples were restricted to PAM50 subtypes LumA, LumB, Her2 and Basal (n=3,184). No sex information was annotated and all samples were used.

*GTEx Data:* GTEx raw count data, sample pheno data and feature data were downloaded using the recount3 (*v1.2.3*) Bioconductor package (Collado-Torres et al., 2017; Wilks et al., 2021). For Machine Learning samples were restricted to those derived from female subjects (n=180). Raw counts were normalized and FPKM transformed using the getRPKM function and values were divided by 2 for paired-end data. FPKM values of transcripts mapping to a gene were summed, and FPKM+0.1 values were log2 transformed consistent with GSE81540 pre-processing.

*The Cancer Genome Atlas Data:* Normalized and pre-processed TCGA RNA-seq FPKM expression values and clinical data from TCGA were downloaded using *TCGAbiolinks* (*v.2.8.4*) (Colaprico et al., 2016) package. RNA-seq expression data were imported using the following parameters: project="TCGA-BRCA"; data.category="Transcriptome Profiling"; data.type="Gene Expression Quantification"; workflow.type="HTSeq - FPKM"; legacy=FALSE. TCGA expression Ensembl IDs were mapped to gene symbols using *EnsDb.Hsapiens.v86*. For visualization and Kruskal-Wallis tests for differences across groups, samples were restricted to those derived from female subjects and samples annotated as either matched normal or PAM50 Normal, LumA, LumB, Her2 and Basal subtypes (n=1,201) and FPKM+1 values were log2 transformed.

For Machine Learning, samples were restricted to matched normal samples and PAM50 subtypes LumA, LumB, Her2 and Basal (n=1,161). PAM50 Normal samples were exluded due to small sample size. FPKM values of transcripts mapping to a gene were summed, and FPKM+0.1 values were log2 transformed consistent with GSE81540 pre-processing.

*SSP OMINER ChIP-Seq Data:* Cistromics ChIP-seq data for each gene of interested was obtained from The Signaling Pathways Project (SPP) Ominer database and restricted to female reproductive system mammary gland data across species. Graphical visualization of ChIP-Atlas MACS binding scores (±10kb from TSS) for *GJB6*, *CLDN10, CLDN11*, *DSC3* and *DSG3* were downloaded from Ominer.

*Functional Annotation of Genes of Interest:* Functional roles of genes highlighted in the manuscript were first explored in The Human Gene Database (*GeneCards v5.0*, https://www.genecards.org/) and subsequent literature search. Transcription factors were identified based on (Lambert et al., 2018).

**QUANTIFICATION AND STATISTICAL ANALYSIS**

All quantification and statistical analyses were conducted using using R (3.5.0) (R Core Team, 2018) (https://www.R-project.org/) and Bioconductor (3.7) (Huber et al., 2015) (https://www.bioconductor.org/) unless otherwise noted.

*Linear regression on expression values between organoids and 4th passage cells:* Transcriptomes in LEPs and MEPs from reduction mammoplasty HMECs at 4$^{th}$ passage (n=11 <30y, n=8 >55y) and uncultured organoids (n=4 <30y, n=3 >55y LEPs; n=3 <30y, n=1 >55y MEPs) were compared using mean subject-level batch-adjusted rlog values calculated for each lineage and age group in organoids and 4$^{th}$ passage cells. Linear regression was performed on the gene expression value means from cells isolated from organoid vs. 4$^{th}$ passage culture in each of the RM LEP <30y, LEP >55y, MEP <30y, MEP >55y subsets. Regression lines and 95% confidence intervals were plotted along with the y-intercept and slope of the line. Linear regression $R^2$ and *p*-value were reported. Residuals were calculated from the linear model (*stats::lm*). Distribution of residuals were plotted, and mean value and sd of the residuals were calculated. Transcripts with absolute value of model residuals ≥ 6 (~4*sd) in either of the 4 subsets were considered outliers.

*Differential Expression Analysis:* For lineage-specific and age-dependent differential expression (DE) analysis, the RNA-seq *edgeR* (*v.3.22.5*) (Robinson et al., 2010) filtered and normalized expression of 17,328 genes (see *DGEList* object in RNA-seq Pre-Processing) were subsetted for $4^{th}$ passage LEP and MEP RM samples (<30y, m=32 LEP and MEP samples, n=11 subjects; >55y, m=22, n=8). DE analysis was performed on sample-level data with linear modeling in *limma* (*v.3.36.5*) (Law et al., 2014; Ritchie et al., 2015) using the *voom* implementation. The experimental design group was defined by the combination of cell type and age group. Batch was modeled into the design matrix (~0 + Design Group + Batch). The four contrast terms included comparisons of MEP vs. LEP lineages in younger <30y and MEP vs. LEP in older >55y groups for lineage-specific DE, and older vs. younger cells in LEP and older vs. younger cells in MEP lineages for age-dependent DE. Sample replicates, as well as the paired nature of MEP/LEP samples were accounted for by calculating the correlation between measurements using the *limma::duplicateCorrelation* function (Smyth et al., 2005) and blocking for subject ID. Because the calculated correlations changed the voom weights slightly, *voom* was re-run for a second time. Correlations were then re-calculated using the new *voom* weights. Linear modeling (*limma::lmFit*) was performed on the *voom* transformed data, with blocking for subject ID. Empirical Bayes moderation (*limma::eBayes*) of computed statistics was then applied.

For lineage-specific DE, contrasts between LEP and MEP in younger <30y and in older >55y women were assesed. Lineage-specific DE threshold was set at Benjamini-Hochberg (BH) adj. $p<0.001$, lfc ≥1 in each age cohort. LEP-specific and MEP-specific genes were defined as those with lineage-specific DE in younger <30y women. MEP-specific and LEP-specific genes were indicated by (+)lfc and (-)lfc respectively. Lineage-specific DE was then annotated as lost, gained or maintained in older women. For age-dependent analyses, contrasts between <30y and >55y LEPs and <30y and >55y MEPs were assessed, and age-dependent directional changes were defined at DE BH adj. $p<0.05$ in each lineage. Genes upregulated in older cells and younger cells were indicated by (+)lfc and (-)lfc respectively.

DE analysis was also performed in *limma* (*v.3.36.5*) (Ritchie et al., 2015) on a publicly available microarray dataset from normal primary breast tissue, GSE102088 (Song et al., 2017).

Normalized $\log_2$ expression data were downloaded from the Gene Expression Omnibus (GEO) using *GEOquery* (*v.2.48.0*) (Davis & Meltzer, 2007). Gene-level data were calculated as the mean expression across all probes mapped to a gene. The experimental design group was defined by age group: younger <30y (n=35), middle-aged >30y <55y (n=68), and older >55y (n=11), in the design matrix (~0 + Design Group). The three contrast terms included pair-wise comparisons between the three age groups. Linear modeling and empirical Bayes moderation of computed statistics with trend parameter=TRUE were applied. Significant age-dependent DE in normal primary tissue was defined as genes with DE adj. $p<0.05$, while nominally significance level was defined as genes with DE unadj. $p<0.05$.

*Differential Variability Analysis:* For differential variability (DV) analysis, RNA-seq count data were batch-adjusted using *ComBatSeq* (*sva_devel*) (Zhang et al., 2020) and bach-adjusted values were normalized using TMM method (*MDSeq v.1.0.5::normalize.counts*); the mean value of *ComBatSeq* batch-adjusted normalized expression data was then taken for subjects with multiple samples. DV analysis was performed using *MDSeq* (*v.1.0.5*) (Ran & Daye, 2017) on subject-level *ComBatSeq* batch-adjusted normalized count data. The experimental design group was defined by the combination of cell type and age group in the design matrix (~Design Group). Outlier removal, which removes outlier genes that are influential upon the effects of covariates, was performed prior to DV analysis with the following parameters: minimum valid sample size threshold was set to 5 samples and significance level cutoff of outlier selections was set to 0.05. Analysis was restricted to the 14,601 genes whose variances could be estimated after outlier removal. The two contrast terms included comparisons of older vs. younger cells in LEP and older vs. younger cells in MEP lineages for age-dependent DV. *MDSeq* was run with default parameters. Testing results were extracted (*MDSeq::extract.ZIMD*) using an inequality test with the following parameters: lfc threshold for inequality testing was set at 0 and with BH *p*-value adjustment method.

For age-dependent analyses, contrasts between <30y and >55y LEPs and <30y and >55y MEPs were performed, and age-dependent variant changes were defined at DV BH adj. $p<0.05$ in each lineage. Genes with increases in variances (higher lfc dispersion) in older cells and younger cells were indicated by (+)lfc and (-)lfc in dispersion respectively.

*Lineage-specific Ligand-Receptor Pair Interactions:* Ligand-receptor pairs (LRPs) (Ramilowski et al., 2015) gene symbols were checked against *EnsDb.Hsapiens.v86* (*v2.99.0*) database (Rainer, 2017) gene symbols and then mapped to Ensembl IDs. Both ligand and receptor median expressions in LEPs and MEPs from younger and older women were calculated from *ComBat* batch-adjusted normalized rlog values. Lineage-specific LRPs were defined based on either the LEP-specific or MEP-specific (DE adj. $p<0.001$ and fold-change ≥2) expression of either the ligand and/or its cognate receptor in the younger cohort. Lineage-specific LRP interactions were considered to be lost in the older cohort when lineage-specific DE of the ligand and/or its cognate receptor was lost in the older cells (not DE at adj. $p<0.001$, lfc≥1). Lineage-specific LRP interactions between ligands and cognate receptors in the younger cohort and lineage-specific LRP interactions lost in the older cohort were visualized in an interactome map (*migest v.1.8.1, circlize v.0.4.8*) (Abel, 2019; Gu et al., 2015). Ligands were connected by chord diagrams from the cell type expressing it (cell type-L-gene symbol) to the cell type expressing its cognate receptor (cell type-R-gene symbol). Functional network enrichment of LRPs in the younger cohort and LRPs lost in the older cohort were performed in STRING database (https://string-db.org/) and enriched KEGG pathways (FDR $p<0.05$) were reported.

*Gene Set Enrichment Analysis (GSEA):* Fast gene set enrichment analysis (*fgsea v.1.6.0*) (Korotkevich et al., 2021) was used to identify age-dependent enrichment of Molecular Signatures Database (MSigDB) (*v7.2*, http://www.gsea-msigdb.org/gsea/msigdb/index.jsp) hallmark gene sets (Liberzon et al., 2015) in LEPs or MEPs. MSigDB gene symbols were mapped to Ensembl

IDs using *org.Hs.eg.db* (*v.3.6.0*) (Carlson, 2018) and mapped to the RNA-seq datasets. Fast GSEA was performed on either rank-ordered DE t-statistics (*limma*) or DV test statistic for dispersion (*MDSeq*) using 1000 permutations. Minimal and maximum gene set sizes were set to 15 and 500 respectively. Enriched gene sets were defined as those with enrichment BH adj. $p<0.05$. For bulk tissue GSEA analysis (GSE102088, <30y n=35, >55y n=11), gene sets were constructed from age-dependent genes in LEPs: (i) 251 genes that were differentially upregulated in young <30y LEPs; and (ii) 220 genes that were that were differentially upregulated in old >55y LEPs. Age-dependent enrichment was assessed in bulk tissue using DE rank-ordered test statistics; enrichment was similarly defined at BH adj. $p<0.05$.

*Protein-protein Interactions and Functional Network Analysis:* Protein-protein interaction (PPI) analysis was performed using the *STRING* database (*v.11*, https://string-db.org/). All possible PPI are considered using all active interaction sources and setting minimum require interaction score to the lowest confidence threshold of 0.150. Network was visualized in *igraph* (v.1.2.5) (Csárdi & Nepusz, 2006), and only the largest fully connected main network of genes was plotted. Community detection was performed on this main network using optimal community structure algorithm in *igraph* (*igraph::cluster_optimal*) (Brandes et al., 2008). Each community was then analyzed in *STRING* for functional network enrichment (FDR $p<0.05$) and common functional terms were summarized and reported.

*Non-parametric Kruskal-Wallis and Post-hoc Wilcoxon Test: Limma*-based genome-wide DE analysis was not performed on publicly available gene expression datasets from breast cancer tissue from the TCGA cohort. Instead, analysis was limited to genes of interest, and non-parametric Kruskal-Wallis (KW) test (*rstatix v.0.5.0::kruskal_test*)(Kassambara, 2020b), a one-way ANOVA on ranks, was used to determine differences in $\log_2$(FPKM+1) values between matched normal and PAM50 subtypes. KW *p*-values were corrected for multiple testing (BH) across all genes when multiple genes were examined. Post-hoc pair-wise comparison between groups was performed using non-parametric Wilcoxon test (*rstatix v.0.5.0::wilcox_test*)(Kassambara, 2020b). Wilcoxon *p*-values were corrected for multiple testing (BH) across all pair-wise comparisons and across all genes when multiple genes were examined. Wilcoxon BH-adj. *p*-values were likewise annotated (*ggpubr v.0.2.5*) (Kassambara, 2020a) at different significance levels ($p$<0.05 (*), <0.01 (**), <0.001 (***), <0.0001 (****).

*Lepage Test on Location and Scale:* Two-sample Lepage test (*nonpar v.0.1-2*) (Pepler, 2017) is a joint non-parametric test of equality for location (central tendency) and scale (variability). Lepage test was performed on the subject-level *ComBat* batch-adjusted normalized rlog expression values of genes encoding for junctional proteins between younger and older cells in each lineage. Significant age-dependent modulation of genes for cell-surface junctional proteins in LEPs and MEPs were defined at $p<0.05$.

*Kolmogorov-Smirnov Test on lineage-specific DE lfc:* Non-parametric two-sample Kolmogorov-Smirnov (KS) test (*stats::ks.test*) on the equality of distributions performed to compare distributions of lineage-specific DE lfc in younger and in older cells. Significance defined at $p<0.05$.

*T-test on the differences of DE lfc between age groups:* One-sided t-test (*stats::t.test*) performed on the distribution of pair-wise differences in lineage-specific DE lfc between age groups (lfc in young - lfc in old) to test if the mean of all values is different from 0. Significance defined at $p<0.05$, nominal significance defined at $p \leq 0.1$.

*T-test on qPCR values between experimental treatments:* Two-sided Student's t-test performed to compare normalized expression between the two groups in co-culture experiments. Significance defined at $p<0.05$.

*Unsupervised hierarchical clustering*: Unsupervised hierarchical clustering in heatmaps (*gplots v.3.0.3::heatmap.2, pheatmap v. 1.0.12*) (Kolde, 2019; Warnes et al., 2020) were implemented using hclust Ward's clustering criterion (ward.D2) agglomerative method with Euclidean distances as distance metric.

*Machine Learning:* Machine Learning gene expression datasets of normal and cancer breast tissue were obtained: GSE81540 (n=3,184) (Brueffer et al., 2020; Brueffer et al., 2018; Dahlgren et al., 2021) GTEx (n=180) and TCGA (n=1,161) as described above. Analysis was restricted to tissues from women annotated as normal or PAM50 LumA, LumB, Her2 and Basal subtypes (PAM50 Normal excluded due to small sample size). Gene features include the 536 of 591 age-dependent DE and DV genes between younger and older LEPs (adj. $p<0.05$) found in common across the GSE81540, GTEx and TCGA datasets. Training and cross-validation was performed on 75% of TCGA data (matched normal=84, PAM50 LumA=425, LumB=156, Her2=62, Basal=146). Test sets for model performance evaluation include 25% of TCGA the model had not seen (matched normal=28, PAM50 LumA=141, LumB=51, Her2=20, Basal=48) and an independent dataset composed of normal tissue from GTEx (n=180) and breast cancer tissue from GSE81540 (PAM50 LumA=1709, LumB=767, Her2=348, Basal=360). Machine Learning was performed using the caret package in R (*v6.0-88*) (Kuhn, 2008) using elastic net implementation from glmnet (*v4.1-2*) (Friedman et al., 2010) using 10-fold cross validation with 3 repeats. To address class imbalance, we used a hybrid subsampling technique implemented using the SMOTE algorithm in the DMwR package (Torgo, 2010). Custom summary functions of model performance were defined, and the model was optimized for mean balanced accuracy which we found to perform better when large class imbalances exist. Model performance in test sets were evaluated using AUC calculated with the MultiROC package (*v1.1.1*) in R (Wei & Wang, 2020) and reported as follows: (i) AUC of each group vs. the rest; (ii) micro-average AUC calculated by stacking all groups together; and (iii) macro-average AUC calculated as the average of all group results. Variable importance values were scaled for each class (range from 0-100%), and gene predictors were identified as genes with scaled variable importance contribution to the predictive model (scaled variable importance >0% in at least one class). Genes with scaled variable importance ≥25% in prediction of at least one class were visualized; gene expression the top 5 predictors in each class were further analyzed in the TCGA breast cancer cohort.
